# Comparative genome analysis of tea rhizosphere dwelling *Paenarthrobacter nicotinovorans* AB444: Uncovering Physiological Resilience and Ecological Adaptability

**DOI:** 10.1101/2025.09.01.673609

**Authors:** Triparna Mukherjee, Sangita Mondal, Dhruba Bhattacharya, Sanjukta Dasgupta, Abhrajyoti Ghosh

## Abstract

Plant growth-promoting rhizobacteria (PGPR) can be considered a promising tool for sustainable agriculture. The recent scenario of growing demand for higher crop yields, with declining soil fertility and impacts of climate change, promotes the global shift towards eco-friendly agricultural practices, emphasising the significance of PGPR as a sustainable solution. In this study, a PGPR *Paenarthrobacter* sp. AB444 was isolated from the rhizosphere of tea plants at the Longview and Makaibari tea estates in the Darjeeling District of West Bengal. Based on the draft genome sequence along with comprehensive chemotaxonomic analyses, strain AB444 was identified as *Paenarthrobacter nicotinovorans*. The pangenomic analysis with phylogenetically close strains revealed the gene clusters that are common and unique to the isolate. The plant growth-promoting attributes of AB444 were explored with different assays like phosphate solubilisation, IAA and siderophore production, ACC deaminase, nitrogen fixation, and protease activity. The colonization of the isolate to the plant root and *in-planta* growth were also assessed. The presence of biosynthetic gene clusters of AB444 related to plant growth was also extensively studied, highlighting the unmapped disposition of *Paenarthrobacter nicotinovorans*.

## 1. Introduction

Rhizosphere is found to be a multifaceted bionetwork where multiple associations are established between microbes and plant root system representing the elementary support system of plant growth [1]. A variety of soil microbes have made a wholesome alliance with the associated plant favouring their growth as well as supporting plants to cope with both abiotic and biotic stress[2,3]. Plant growth-promoting rhizobacteria (PGPR) are a substantial microbial community associated symbiotically with plant roots, supporting plant growth using various processes like nitrogen fixation, plant hormone production (cytokinin, auxins, and gibberellins), phosphate solubilization, production of siderophores, and plant senescence inhibition[4]. Beneficial effects of these PGPR on plants include improved root and shoot growth, their biomass, nutrient uptake efficiency, chlorophyll content, total protein content, abiotic stress tolerance, and hydraulic activity with deferred senescence[4]. Some other indirect means of plant growth promotion include the production of exopolysaccharides, hydrolytic enzymes, heavy metal bioremediation, along with induced systemic resistance (ISR)[5,6]. Certain PGPR exhibit diverse chemotypical patterns, contributing to their effective antifungal activity as biocontrol agents. This considers the production of antimicrobials, siderophores, lytic enzymes with multiple extracellular metabolites that will intervene with the growth of the broad microbial host range [7]. PGPR can overturn disorders by promptly synthesizing pathogen-alienating compounds and by stimulating the immune responses of the plant. PGPR triggers the plant immune response with the initiation of induced systemic resistance by means of establishing plant responses to environmental strain in relation to the physical as well as biochemical parameters [6,8]. PGPR, in association with host systems, are often found to enhance the biosynthesis of bioactive defence-aiding molecules. These molecules assist the host plant in better surviving stressful conditions by exerting direct effects against phytopathogens[8]. The antimicrobial responses accompanied by the synthesis of a variety of plant growth promoters (siderophores, hormones), suggest that the PGPR could be explored for dual-purpose approaches on the basis of the application of a particular preparation working as both biofertilizer and biopesticide also [9]. In this regard, microbial populations like filamentous actinomycetes create an exceptional hold among multiple microbial occupants for supporting the plant survival.

PGPR are highly diverse, belonging to various suborders, including sub-order Micrococcineae, with considerable variation across genera. A well-known genus within the Micrococcaceae family is *Arthrobacter*, comprising aerobic bacteria that exhibit coccoid or coryneform morphology. *Arthrobacter* spp. show immense metabolic diversity with the ability to produce a number of valuable metabolites like amino acids, enzymes, vitamins, and polysaccharides, specific growth factors[10,11]. *Arthrobacter* spp. are mostly valued for their role in plant growth-promoting activities; however, their potential as contributors to secondary metabolites remains underexplored. Recently there have been some reports regarding the production of bioactive metabolites by some *Arthrobacter* spp.[12]. Until 2016, the genus *Arthrobacter* were categorized in many species that were classified sequentially into 5 new genera, counting *Paenarthrobacter* as one of them[13].

*Paenarthrobacter* adv. *Paene*, of that is the conversion from Latin word is “almost” or “closely” (Glare, 1968) (nearly Arthrobacter), is a Gram-positive bacterial genus reclassified to include six species: *Paenarthrobacter nicotinovorans, Paenarthrobacter aurescens, Paenarthrobacter ilicis, Paenarthrobacter histidinolovorans*, *Paenarthrobacter nitroguajacolicus*, and *Paenarthrobacter ureafaciens*[14]. However, the metabolic patterns and functionalities of *Paenarthrobacter* remain to be precisely explored. The extensive use of biofertilizer to comply with the global need has driven efforts to discover potential multifaceted bioinoculants in unexplored regions, including the North-Eastern segment of India, with a particular emphasis on the tea (*Camellia sinensis* (L.) O. Kuntze) rhizosphere. Tea is considered as commercially valued perennial crop generally planted in North-Eastern segment of India. However, studies on the Indian tea rhizosphere microbiome remain limited, revealing the presence of *Arthrobacter* spp. [5,6].

In this context, our study aims to isolate and characterize *Paenarthrobacter* species from the tea rhizosphere of Darjeeling, with a particular focus on its potential role in plant nutrition and mitigation of biotic stress through the metabolites produced by it. Comprehensive genomic studies, coupled with the identification of biosynthetic gene clusters (BGCs) in *P. nicotinovorans* AB444, will provide insights into the genetic basis for its role as a biotic stress-reliever in the case of plant growth-promoting activities. Here, this study focuses on the anti-phytopathogenic activity of the genus Paenarthrobacter reflecting its antimicrobial potential, which comes as a rarely explored functionality of Paenarthrobacter till date. The comparative phylogenomic approach will underscore the evolutionary relationships between *P. nicotinovorans* AB444 and closely related species, highlighting unique genomic traits that contribute to its efficacy as a potent PGPR. The findings are expected to contribute significantly to the development of sustainable agricultural practices by harnessing the untapped potential of *Paenarthrobacter* species in enhancing crop productivity and resilience in tea plantations and beyond.

## 2. Materials and methods

### 2.1 Isolation of Actinobacteria

Rhizospheric and bulk soil samples were collected from the tea plants of two estates: Longview tea estate and Makaibari tea estate of Darjeeling District of West Bengal. Sampling was carried out from 20 ± 2 cm depth of tea rhizospheric zone. Each tea estate was divided into three distinct sections to minimize sampling bias. From each section, samples were randomly collected from four healthy tea plants, including their roots and the surrounding root-adhering soil. The four rhizospheric soil samples from each section were combined, resulting in three composite rhizospheric soil samples from each tea estate, prepared for downstream analysis. Soil samples were compiled in sterilized dry polythene bags using sanitized gloves to avoid any microbial contamination. Collected soils were preserved at 4 °C until further processed. The latitude and Longitude were recorded by using GPS tracking gadget (Table 1). The physicochemical characters of collected soil samples were analysed following the protocol given by[15].

#### 2.1.1 Bacterial strain, growth condition, and culture maintenance

Saline water (0.9 %) was used to make suspension of the collected soil samples and serially diluted upto 10^−6^. Subsequently, 100 μl of the diluted samples were spread onto two different selective media. The compositions of the media were as follows-M2 Media (g/l): Sodium caseinate-2 g; L-asparagine-0.1 g; sodium propionate-4g; K_2_HPO-0.5 g; MgSO_4_, 7H_2_O-0.1 g; FeSO_4_,7H_2_0-0.01 g; agar-12 g; (pH 8.1 ± 0.2). ISP2 Media (g/l): Yeast extract-4 g; malt extract-10 g; Dextrose-4 g; agar-12 g (pH 7.2 ± 0.2)

#### 2.1.2 Microscopic Identification

The phenotypic property of the bacterial isolate AB444 was investigated as per the standard microbiological protocol. Particular actinobacterial isolates were initially lay open to phase contrast microscope (with Zeiss Scope A1 microscope) to notice the positioning of branched aerial hyphae. The morphological pattern of the vegetative cell was also explored from scanning electron microscopic (SEM) study. Here log phase bacterial cells were fixed in 0.25% glutaraldehyde followed by subsequent dehydration by gradual ethanol concentration. Using sputter, the gold coating of the bacterial sample was done and visualised under an electron high tension (EHT) of 15kV (ZEISS EVO18) with thick gold coating in Ruorum 150 TES for 10 min.

#### 2.1.3 Biochemical characterization based on carbon utilization

The GEN III MicroPlate™ test, a standardized Biolog method using 94 biochemical tests, was done with the selected strain to get a profile and identify a broad range of Gram-positive and Gram-negative bacteria[16]. The biolog GEN III MicroPlate™ studies of the microbe based on phenotypic tests included 71 carbon source utilization assays and 23 chemical sensitivity assays. All necessary nutrients and biochemicals are prefilled and dried into the 96 wells of the MicroPlate. The tetrazolium redox dye was employed as a colorimetric indicator to assess the utilization of carbon sources or resistance to inhibitory chemicals.

#### 2.1.4 Chemotaxonomic identification

The Fatty Acid Methyl Ester (FAME) analysis was conducted to achieve the chemotaxonomic characterization of the isolate. This analysis was outsourced to Royal Life Sciences Pvt. Ltd., an ISO 9001:2015 certified organization affiliated with MIDI Sherlock, USA, India. The study was done through typical gas chromatography with applying then MIDI Sherlock MIDI Sherlock system.

### 2.2 Plant growth promoting attributes

#### 2.2.1 Phosphate solubilization

The ability of AB444 to solubilize insoluble phosphate was screened by both qualitative and quantitative methods. Total 2µl of freshly grown bacterial culture was spotted on Pikovsaya’s agar plate supplemented with insoluble tricalcium phosphate and incubated at 28°C for 7days [17]. Appearance of clear halo zone was considered as positive result and the solubilization index (SI) was calculated.

Solubilization index (SI) = (Zone diameter/ Colony diameter)

Similarly, quantitative evaluation was done according to the method of [17]with certain modifications. 50µl of freshly grown bacterial culture was inoculated in 5ml of Pikovskaya’s Broth supplemented with insoluble tricalcium phosphate. This culture was incubated at 28°C under shaking conditions (at 146rpm) for seven days. After 7days, supernatant containing the soluble phosphate was collected by centrifugation at 10,000rpm for 20mins. Quantification of soluble phosphate was done using Chen reagent (6N H_2_SO_4_, sterile H_2_O, 1%ammonium molybdate tetrahydrate and 10% ascorbic acid at 1:2:1:1 ratio). 1ml of the supernatant was mixed well with 4ml of Chen reagent followed by incubation at 37°C for 1 hour and 30 minutes [6]). Uninoculated media was considered as control. Formation of the blue-coloured ammonium-phosphomolybdate was measured spectrophotometrically at 619nm (Halo XB-10 UV-vis single beam spectrophotometer by Dynamica Scientific Lmt, UK) after diluting it with 3ml of sterile water. The amount of phosphate solubilization was calculated from the standard curve of KH_2_PO_4._

#### 2.2.2 Indole acetic acid production

From a freshly grown culture, 50µl was inoculated in 5ml Luria-Bertani broth either supplemented with or without tryptophan and incubated at 28°C with continuous shaking. Finally, after 7 days of incubation, the cultures were centrifuged at 10,000 rpm for 10 minutes to pellet the cells. The resulting cell-free supernatant was thoroughly mixed with Salkowski reagent in a 1:2 ratio, followed by incubation in the dark at room temperature for 30 minutes. The intensity of IAA complex formation was measured spectrophotometrically at 535nm (Dynamica XB-10). Uninoculated media with or without tryptophan was considered as control. IAA production was calculated from the standard curve of commercial IAA (Sigma-Aldrich Germany)[18].

#### 2.2.3 Siderophore production

The ability to produce siderophore was screened by using chrome azurol S method [19].CAS agar plate was inoculated with 2µl of freshly grown bacterial culture and incubated at 28°C for 7days. The appearance of an orange halo was considered as a positive result. Quantification of the siderophore was done using the CAS-shuttle solution[20]. The blue dye was prepared with certain modifications. MM9 media was inoculated with 1% of bacterial culture and incubated at 28°C at 146rpm. After 7days post-incubation, supernatant was collected by centrifugation at 3,000 rpm for 15 minutes. A volume of 500 µL of cell-free supernatant was combined with 500 µL of CAS-shuttle solution, followed by the addition of 10 µL of 3% sulfosalicylic acid. The mixture was incubated in the dark at room temperature for 20 minutes. Siderophore production was quantified spectrophotometrically at 630 nm, using an uninoculated control processed in the same manner as the reference sample.

% of siderophore production _=_ [(A_r_-A_s_)/A_r_]*100

Where, A_r_ is the absorbance of reference and A_s_ is the absorbance of the sample at 630 nm.

#### 2.2.4 Protease activity

To screen for its protease activity, 2µl volume of freshly grown culture was spotted on skim milk agar (SM agar) and incubated at 28°C for 7 days. The appearance of halo zone was determined as a positive result [21].

#### 2.2.5 ACC deaminase production

For screening of ACC deaminase production ability, the isolate was inoculated on minimal media agar plate supplemented with 3mM ACC (1-aminocyclopropane 1-carboxylic acid) as sole nitrogen source [5]. Plates were incubated at 28°C for 7 days. Appearance of growth was considered as a positive result.

#### 2.2.6 Nitrogen fixation

The isolate was separately grown on nitrogen-free Burk’s medium to screen for it’s nitrogen fixation ability. Plates were incubated at 28°C for 7 days. Appearance of growth was considered as positive result[22].

#### 2.2.7 Plant growth promotion by root colonization

*In-planta* growth promotion experiment was done by inoculating the isolate onto maize plants. Surface-sterilized maize seeds were sown on sterile soil-rite (Satya Agro Irrigation Co., West Bengal, India). The inoculum was prepared by inoculating 10ml ISP2 media with the glycerol stock of AB444 and incubated at 28°C and 146rpm. On the day of treatment, culture was pelleted down at 0.6 (O.D_600_) by centrifugation at 6,000rpm for 15mins. The resulting pellet was carefully resuspended in 5ml of 20mM potassium-phosphate buffer (pH 7). After germination, each seedling was inoculated with the 500µl of prepared inoculum and only buffer was used as the control. This treatment was done twice at 7 days of interval up-to 14days. Finally, plants were harvested at 21^st^ day. Plants were carefully detached from the soil-rite and roots were washed gently under tap water. Both root and shoot lengths were measured. Fresh weight and dry weight of treated and control plants were documented.

For the colonization evidence, fresh roots of control and treated plants were cut into small pieces and washed twice with 1X PBS (pH 7) with gentle care. The sample fixed in 1% glutaraldehyde solution dissolved in 1X PBS for 1 hour in dark at room temperature. The fixed root samples were washed again with 1X PBS buffer. The sample was dehydrated by incubating at different alcohol gradations (30%, 50%, 70%, 80%, 90%, 95%) for 15mins each. Final dehydration step included incubation at 100% ethanol. The sample was placed on grease-free glass slide and gold-coated for visualization under a scanning electron microscope (SEM) [FEI Quanta 200 MK] [6].

### 2.3 Antimicrobial activity of AB444

Antimicrobial activity of the crude extract of AB444 was screened against two phytopathogens, vis. *Ustilago maydis* SG200 and *Xanthomonas campestris pv. campestris*. Primary culture was prepared by inoculating YEPS broth with the former strain whereas PSB broth was inoculated from glycerol stock of the later one. Cultures were incubated overnight at 28°C and 146rpm. Freshly grown culture was diluted at 1:100 ratio. In a 96 well plate, crude extract of AB44 was added to 200µl of the diluted culture as required to maintain 400µg/ml, 500µg/ml, 600µg/ml, 700µg/ml, 800µg/ml concentrations. Untreated culture was considered as the negative control, whereas sterile media and commercial antibiotic treated culture was considered as positive controls. Cycloheximide (50µg/ml) and kanamycin (50µg/ml) were used against yeast and bacteria, respectively. Plate was incubated overnight at 28°C and 146rpm. After the appearance of prominent growth in negative control, growth was observed by measuring the absorbance at O.D_600_ in a plate reader (Thermo Multiscan Go cat no. 51119300). Cell viability was observed by using resazurin dye. To visually assess the cell viability, 2µl of 5mg/ml resazurin dye was added to each sample and the plate was incubated at room temperature for 30mins. Appearance of pink colour represented cell viability whereas purple colour was considered as cell death. Each set was performed in triplicates.

### 2.4 Isolation of genomic DNA and draft genome sequencing

The genomic DNA was extracted from AB444 cultures using NucleoSpin Microbial DNA kit (Macehery-Nagel, Germany) and the draft genome sequencing was performed on Illumina MiSeq platform [23]. The initial quality assessment of the paired end raw fastq reads of the sample was performed using FastQC v.0.11.9 with default parameters [24]. Thereafter, quality checked raw fastq reads were trimmed and processed using Fastp v.0.20. [25], finally quality re-assessment was done using FastQC. The processed paired-end reads were assembled using Unicycler [26]. Thereafter, the completeness and contamination of the genome were checked with DFAST [27]. The 16S rRNA gene fragment from the assembled genomic FASTA was extracted using the ContEST16S [28] tool of EzBioCloud [29]. The similarity-based search of AB444 genome was performed using EzBioCloud’s ‘16S based ID’ and the top-hit taxon was identified. The Average nucleotide (ANIb) identity was assessed at JSpeciesWS [https://jspecies.ribohost.com/jspeciesws] and the species of the isolate was determined by calculating DNA-DNA hybridization (DDH) [<u>ggdc.dsmz.de</u>] among the available *Paenarthrobacter nicotinovorans* genomes from NCBI. The raw sequence reads and assembled draft genome were submitted in NCBI under the accession number PRJNA931222.The circular map of AB444 was visualised with Proksee web-based tool [30]. After assembly, the AB44 genome along with 19 *P. nicotinovorans* genomes obtained from NCBI were annotated with Prokka version 1.14.6. [31].

#### 2.4.1 Comparative genome analysis

Pangenome analysis was subsequently carried out using Roary version 3.13.0 [32] with the minimum percentage identity for BLASTp set to 80%. The core-gene phylogeny was inferred from the single copy core-gene alignment generated from Roary. A maximum-likelihood tree was reconstructed from aligned single copy core-gene using RAxML version 8.2.12 [33]. The gene cluster output from Roary were further annotated using the Kyoto Encyclopedia of Genes and Genomes (KEGG) database [34].

#### 2.4.2 Exploring biosynthetic gene clusters

Antismash was used to get information regarding biosynthetic gene cluster of AB444 and phylogenetically closed species. A heatmap was generated using the Clustering Visualizer (Clustvis) web tool to visualize the correlation between genes and their corresponding genera. Additionally, an unsupervised machine learning model, Principal Component Analysis (PCA), was developed to analyze the data. To enhance the discriminatory capabilities of the model, a supervised Partial Least Squares Discriminant Analysis (PLS-DA) model was also constructed. This analysis was conducted using the R Bioconductor package. Overall, these methods allowed for a comprehensive exploration of the relationships between genes and genera, with the integration of both unsupervised and supervised machine learning techniques to uncover meaningful patterns in the data.

## 3. Results

The present work involved a comprehensive morphological and biochemical characterization of the AB444 strain, identified as a plant growth-promoting bacterium and isolated using selective actinobacteria-specific media. The cultures of AB444 showed slimy beige colour colonies on M2 media and in the presence of long light exposure, the colony colour changed from beige to yellowish (Figure S1). Phase contrast micrographs of AB444 cells were found as rod shaped, cells are arranged in regular chain/clusters, tetrads, as well as in pairs with some cell forming V-shaped arrangement. Typically, *Paenarthrobacter* species are known to undergo a rod-coccus growth cycle, appearing as rods during the exponential phase and transitioning to cocci in the stationary phase. In this study, a small number of round-shaped cells were noted, potentially indicating the transition phase of the growth cycle. Scanning electron micrographs of strain AB444 revealed a rod-shaped cellular morphology with a smooth surface exhibiting a gelatinous texture, consistent with observations made using phase-contrast microscopy. Cells of AB444 were found to be present either single or in a group or both (Figure S2). From the Biolog GEN III MicroPlate analysis comprehensive information regarding the biochemical pattern has been characterized (Table S1). Summarized result analysis connects to the phenotypic fingerprinting along with the genus-level confirmation. The outcome of the fatty acid methyl ester (FAME) study is presented in supplementary file 1: Table S2. The predominant cellular fatty acids of AB444 were identified as 14:0 ISO (2.88%), 15:1 ANTEISO A 7.23 %), 15:0 ISO (5.33 %), 5:0 ANTEISO (49.07 %), 16:0 ISO (19.69%), 16:0 (2.43%), 17:0 ANTEISO (10.87%).

The isolate AB444 showed varied range of PGP activities. After 7 days of incubation the isolate showed clear halo zone in Pikovskaya’s agar (SI =1.5). From quantitative estimation, we found that it can solubilize 117.66 ± 0.5 µg/ml of inorganic phosphate. The isolate is capable of producing siderophore which can be shown from the orange halo on CAS agar plate (Figure S3). The percentage of siderophore produced by this strain is 13.46 ± 0.2. It also showed clear halo zone in SM agar (Figure S3) ensuring its ability to break down proteinaceous substances, might be able to inhibit pathogens. It showed moderate growth on ACC supplemented M9 agar plates indicating positive result for ACC deaminase production (Figure S3). But it showed no growth on Burk’s medium indicating it might not have the potential for nitrogen fixation. This isolate showed gradual increase in IAA production (Figure S3), although it can produce even in the absence of the IAA precursor i.e tryptophan which indicates that it has both tryptophan dependent and independent indole pathway.

### 3.1 In-planta growth promotion by root colonization

AB444 inoculated plants showed better root length than the uninoculated control plants, whereas shoot length was very less altered in the inoculated plant (Figure 1). Root association is an important function of any PGPR. It ensures physical attachment of the rhizobacteria with the plant establishing a strong interaction with the respective host plant. SEM images clearly showed that this isolate is capable of colonizing in roots of maize plants (Figure 2). The root colonizing ability of AB444 suggests that it could be a potent growth-promoting rhizobacterium.

**Figure 1:**
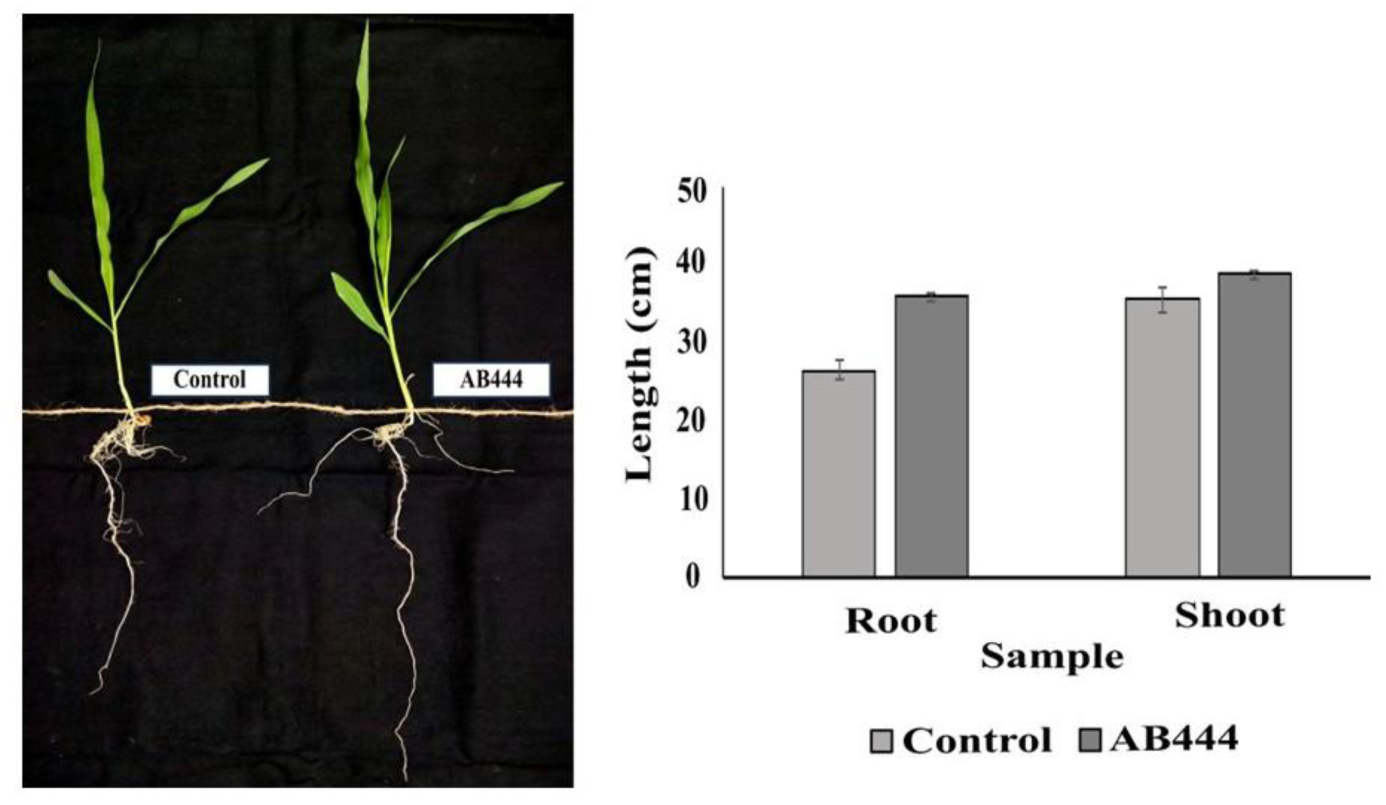
Plant growth-promoting trait of AB444

**Figure 2:**
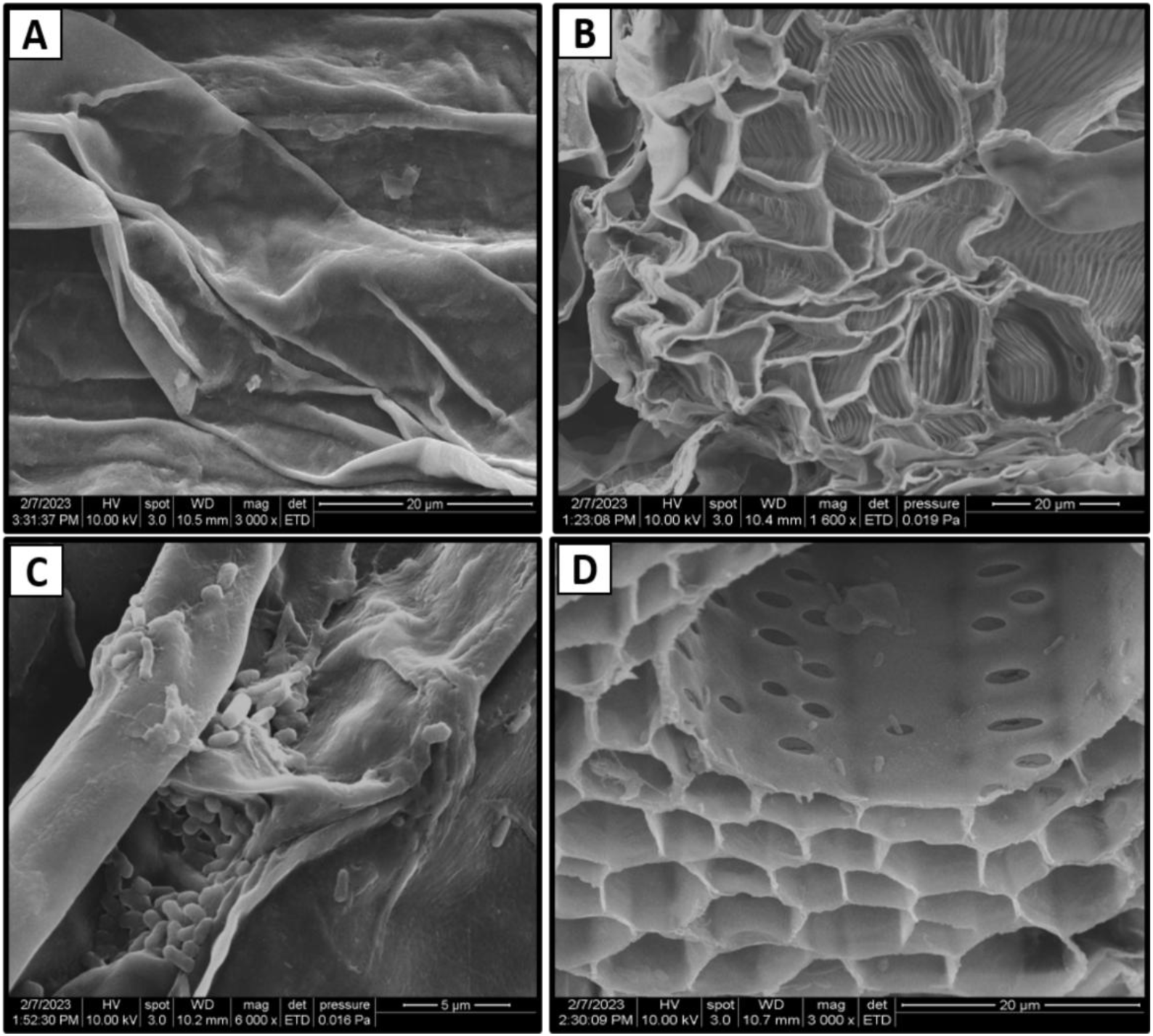
SEM micrograph of root colonization of AB444 strain. A and B. Uninoculated control root. C and D. Roots inoculated with AB444

### 3.2 Phytopathogenic activity of AB444

We tried to find out phytopathogenic activity of AB444 strain against two harmful pathogens of maize plant, vis. *Ustilago maydis* SG200 and *Xanthomonas campestris pv. campestris* strains. *U. maydis* SG200, a fungal phytopathogen causes smut disease by forming galls on the maize shoots. Whereas *Xanthomonas campestris* is a gram-negative bacterial pathogen of cruciferous plants causes blight diseases. AB444 strain was potent in inhibiting the growth of both *Xanthomonas* sp. and *Ustilago maydis* under *in-vitro* conditions. The growth of *Xanthomonas* sp. was inhibited at 500µg/ml of crude extract of AB444 whereas, SG200 growth was effectively reduced at 400µg/ml concentration (Figure S4).

### 3.3 Taxonomic affiliation of AB444

The BLAST search and EzBioCloud analysis of 16S rRNA gene obtained from AB444 draft genome sequence revealed the highest homology with *Paenarthrobacter nicotinovorans* DSM 420 (99.58%) followed by *Paenarthrobacter histidinolovorans* DSM 20115 (99.30%), and *Paenarthrobacter ureafaciens* DSM 20126 (98.88%) (Table S2). Therefore, the genus of A444 was identified as *Paenarthrobacter* sp. Total of 18 available genomes of *Paenarthrobacter nicotinovorans* were selected from NCBI for ANIb, DDH and pangenomic analysis.

### 3.4 Results of genome analysis

The genome assembly of strain AB444 resulted in 42 contigs, comprising a total of 3,988 genes. Among these, 3,927 are coding sequences (CDSs), with 3,904 genes coding for proteins. The genome also contains 61 RNA genes, including one complete 16S rRNA and two partial 23S rRNAs (Figure S5). Additionally, the genome encodes 55 tRNAs and three non-coding RNAs (ncRNAs). This comprehensive annotation highlights the genetic potential of AB444, supporting its diverse functional capabilities. The ANIb result of AB444 along with 18 selected *Paenarthrobacter nicotinovorans* genomes revealed that the highest ANI% of AB444 was observed against *P. nicotinovorans* NPDC089694 (98.35%) (Table S3, Figure 3). Whereas, the maximum DDH% was observed with *P. nicotinovorans* NPDC089692 (90.8%) and *P. nicotinovorans* NPDC089694 (90.8%) (Table S3). The pangenomic analysis showed total of 17494 genes out of which 900 and 80 were identified as core and soft-core genes, respectively whereas total of 8500 shell genes and 8014 cloud genes was estimated (Figure S6). A phylogenetic tree of *Paenarthrobacter nicotinovorans* strains including AB444 was constructed to analyse their genetic diversity and relatedness. The phylogenetic tree constructed demonstrates the close relatedness of the AB444 genome with the genomes of *Paenarthrobacter nicotinovorans* NPDC089692, *Paenarthrobacter nicotinovorans* DS1097, *Paenarthrobacter nicotinovorans* NPDC089711 and *Paenarthrobacter nicotinovorans* NPDC089708. The tree is well-supported, with bootstrap values exceeding 80% for most nodes, indicating reliable clustering. This finding was further supported by ANIb and DDH values. The heatmap generated from pangenomic analysis (Figure 4) illustrated the presence and absence of specific genomic features among the strains. Distinct patterns of genomic variation were observed, highlighting genetic differences between closely related strains. Certain genomic features are consistently present in specific clades, suggesting the functional conservation within these groups. Conversely, distributions of other features are not uniformly distributed, reflecting potential horizontal gene transfer or gene loss events. Notably, closely related strains such as *Paenarthobacter* AB444 and *P. nicotinovorans* NPDC089708 exhibit unique genomic profiles that distinguish them from other strains.

**Figure 3:**
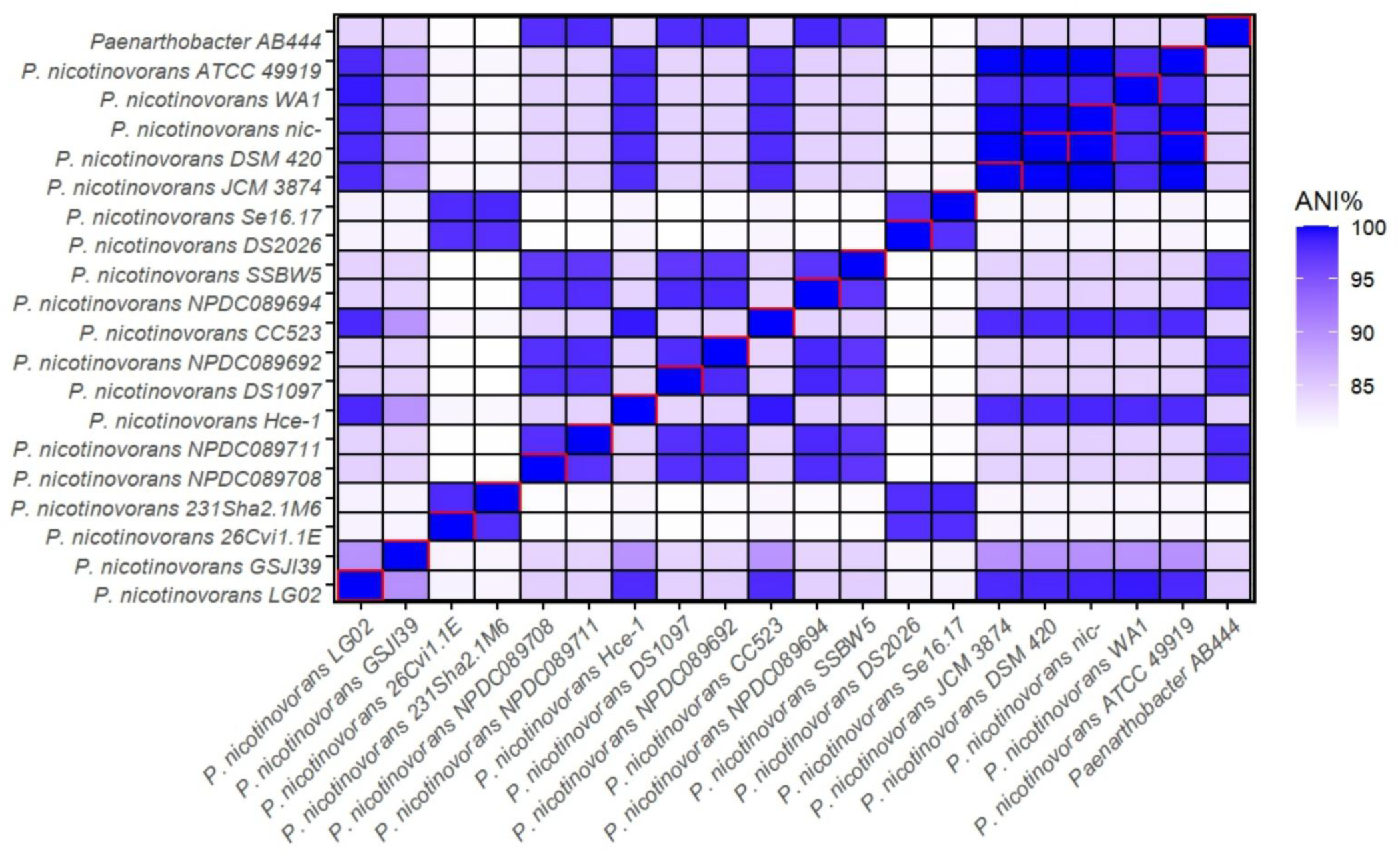
Average nucleotide identity (ANI) matrix of AB444 and *Paenarthobacter nicotinovorans* strains

**Figure 4:**
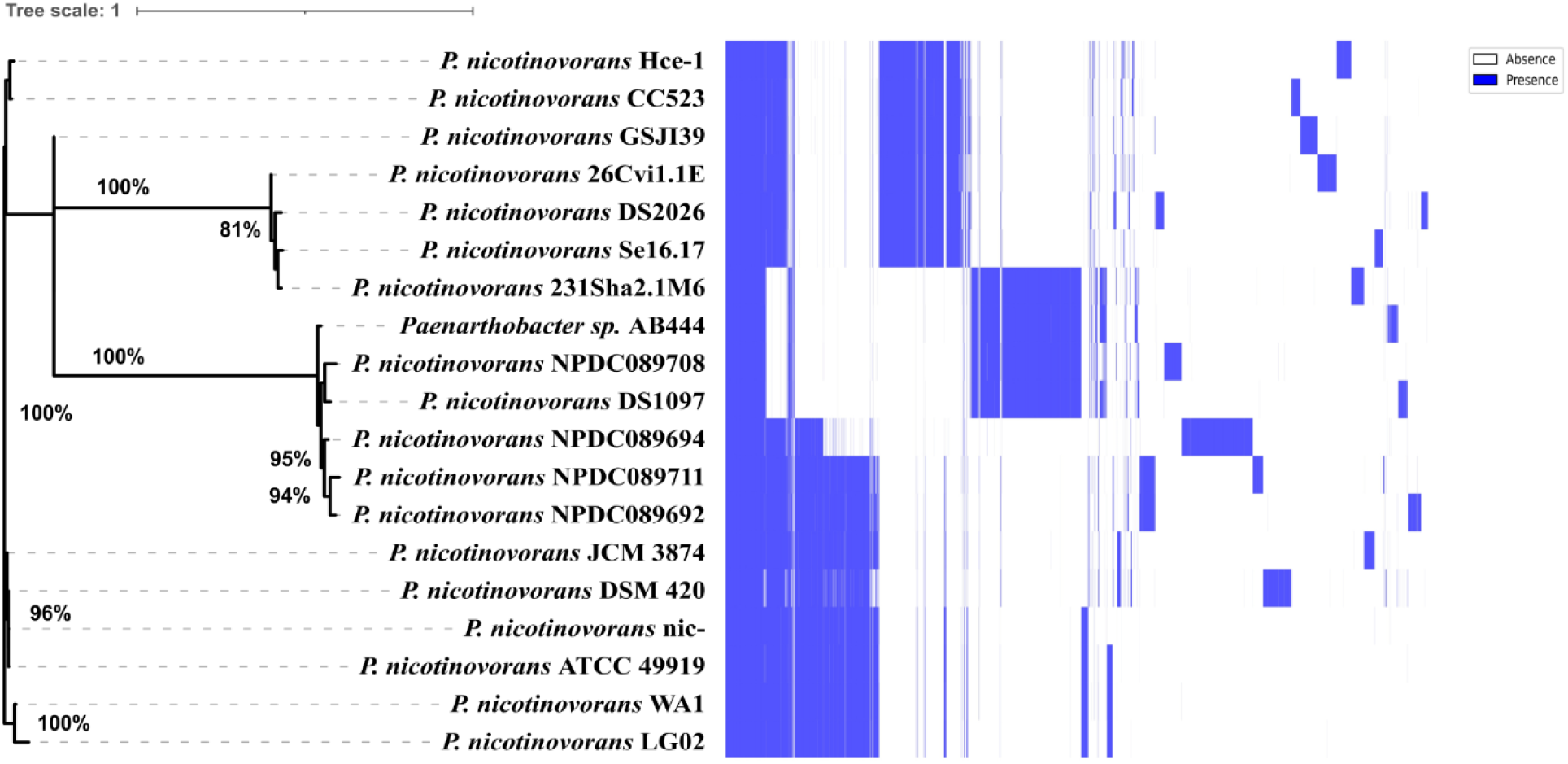
The phylogenetic tree (left) and pangenome gene present-absent matrix of AB444 and *Paenarthobacter nicotinovorans* strains

The KEGG functional annotation of AB444 and *P. nicotinovorans* strains categorized the gene clusters into core, accessory, and unique genes (Figure 5). The core gene clusters are conserved across all strains and associated with essential metabolic functions, including amino acid metabolism, nucleotide metabolism, and carbohydrate metabolism. These clusters likely to play a crucial role in maintaining basic cellular functions and viability. Accessory genes among the strains were predominantly associated with pathways such as membrane transport, xenobiotics biodegradation and metabolism, and signal transduction. These genes may provide adaptive advantages, enabling strains to survive in diverse environmental conditions. Additionally, unique gene clusters, were mainly linked to specialized functions, including biosynthesis of secondary metabolites, environmental adaptation, and drug resistance. These genes reflect strain-specific traits, highlighting their potential ecological roles or interactions with specific niches. Notably, a large number of gene clusters were mapped to metabolism of cofactors and vitamins, suggesting the importance of these pathways in the metabolic versatility of AB444 and *P. nicotinovorans* strains. Similarly, pathways associated with environmental information processing and xenobiotics biodegradation suggest the potential of these strains for bioremediation.

**Figure 5:**
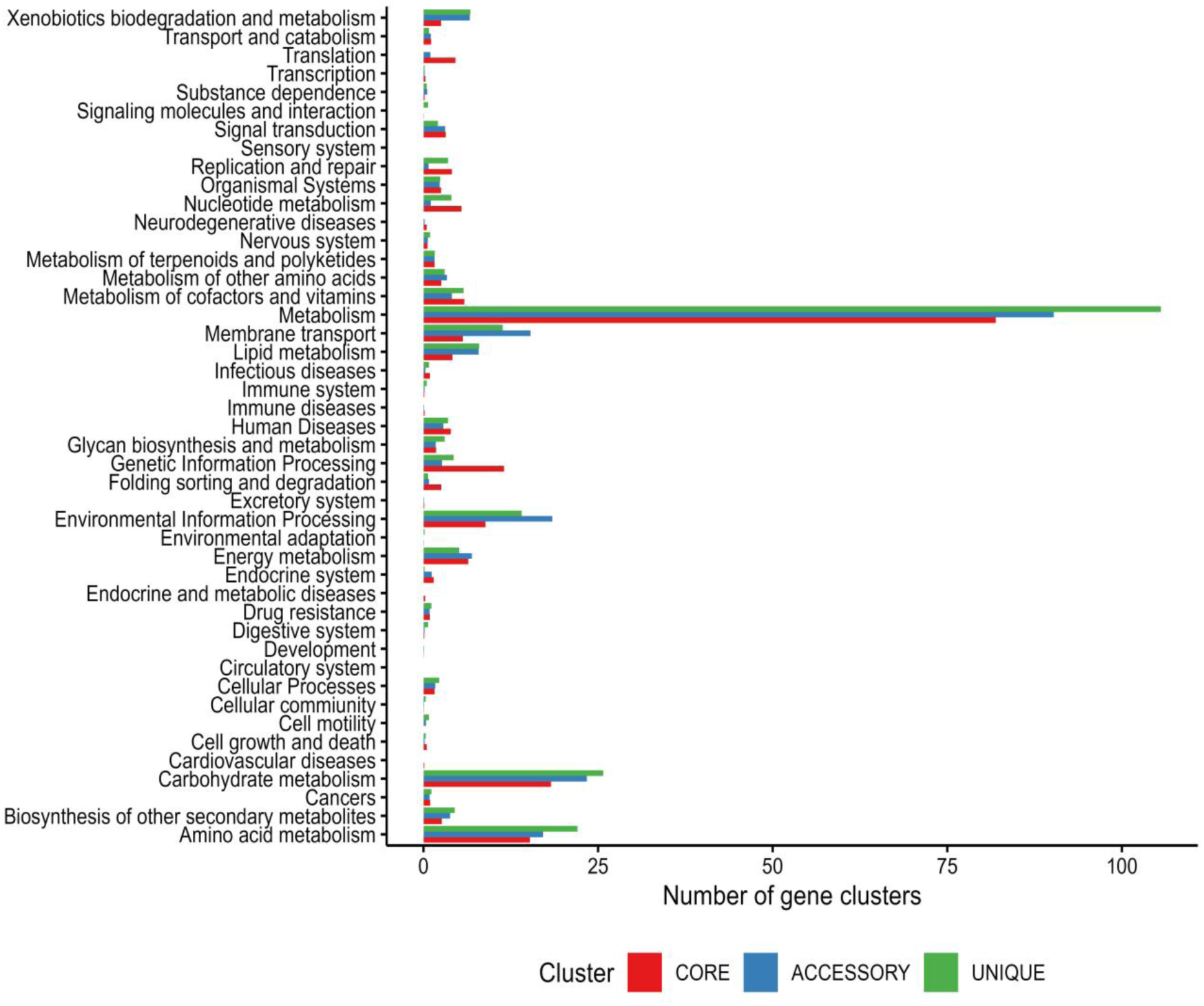
The KEGG annotations of the AB444 and *Paenarthobacter nicotinovorans* strains pangenome

### 3.5 Results of biosynthetic gene cluster

The genome analysis of *Paenarthrobacter* sp. in antiSMASH (“relaxed” strictness) disclosed 6 regions of biosynthetic gene clusters. Figure 6 presents a metabolite to organism correlation heatmap illustrating the correlation between species and gene cluster, with correlation values ranging from -2 to +5. Negative correlations are denoted by blue, while positive correlations are represented by red. In the case of *Paenarthrobacter* AB444 major corelation was found with the Other:Cyclitol gene cluster and then followed by terpene, other, NRP Cyclic depsipeptide, polyketide. *Arthrobacter* sp. GN70 also shows more close association with Other:Cyclitol gene cluster and it shows more likely relation with *Paenarthrobacter* AB444. Metabolite-to-metabolite correlation heatmap is shown in Figure 7. The correlation ranges from -0.7 to +1.0. The red boxes in the heatmap denote the negative correlation, whereas the blue ones show the positive associations. A diverse response was found and established a fundamental relationship pattern of biosynthetic gene cluster among selected organisms. Additionally, a dendrogram plot is included to visualize the hierarchical clustering between the groups based on the biosynthetic genes (Figure 8a) and a closer association was found with *Arthrobacter* sp.

**Figure 6:**
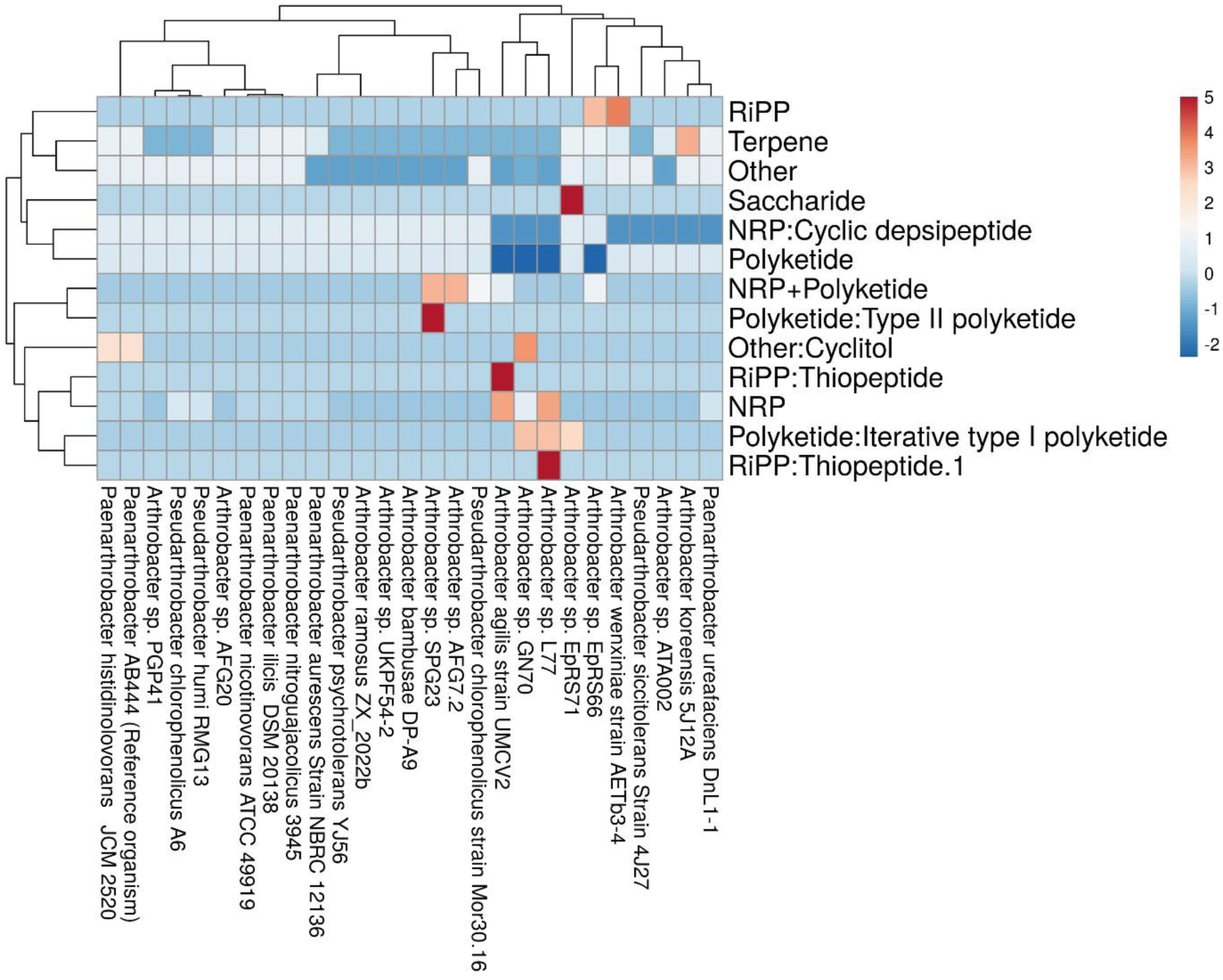
The metabolite-to-organism correlation heatmap

**Figure 7:**
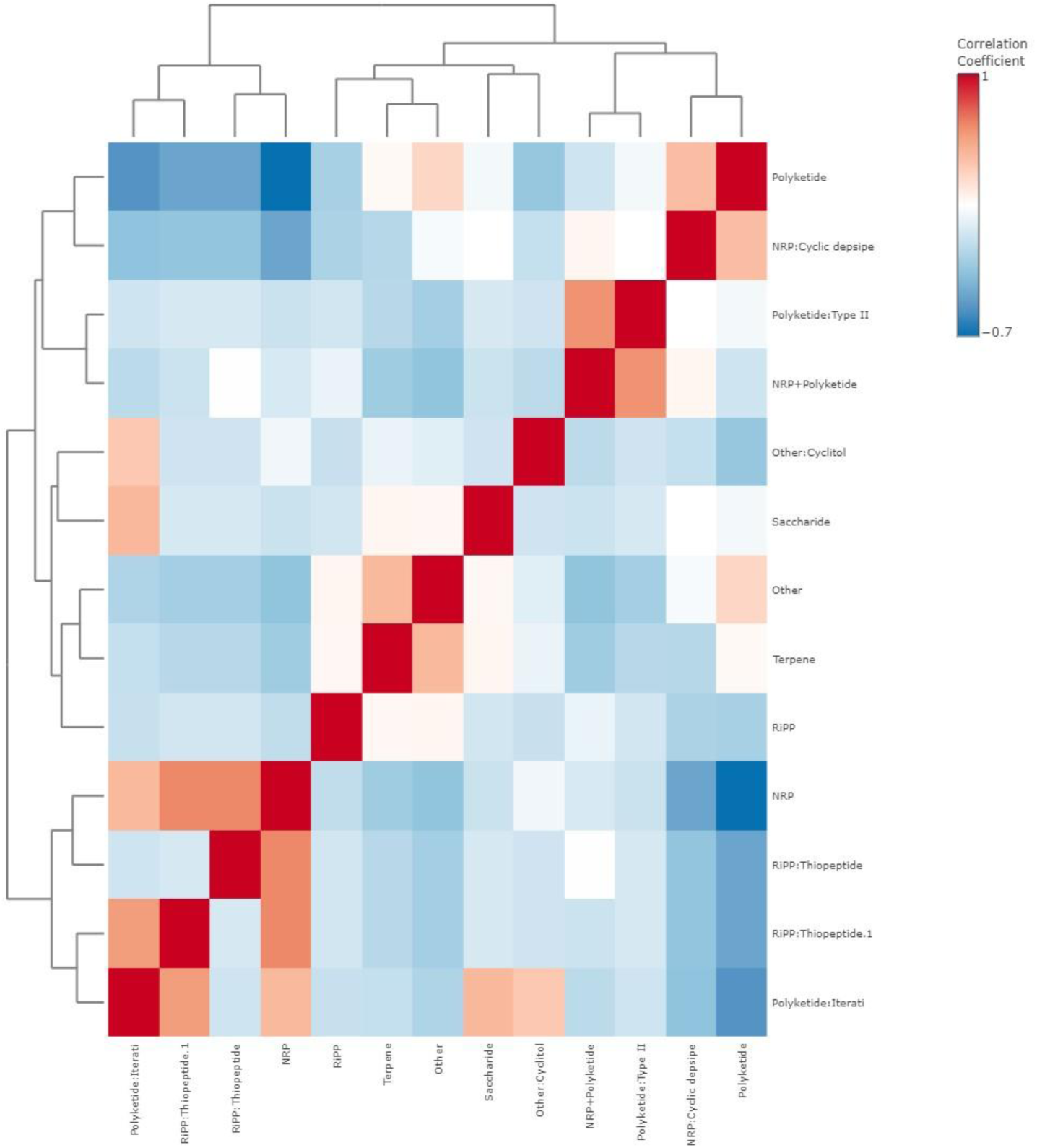
Metabolite to metabolite heatmap based on their expressions in the selected organism’s dendrogram

**Figure 8.**
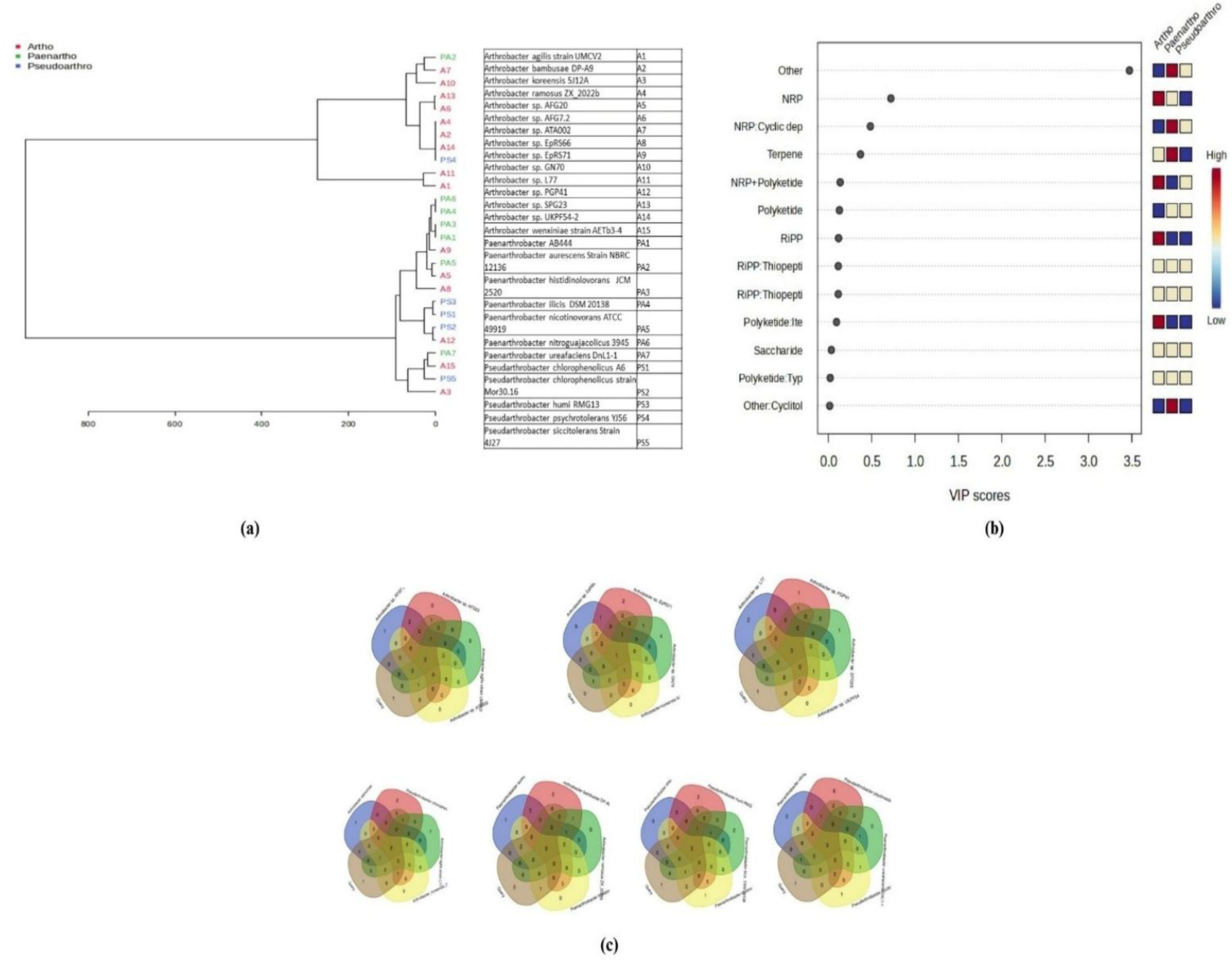
**(a):** A dendrogram plot is included to visualize the hierarchical clustering between the groups based on the biosynthetic genes, **(b):** VIP plot discriminating among the *Paenarthrobacter* sp*, Pseudarthrobacter* sp and *Arthrobacter* sp , **(c):** Venn Diagram of overlapping genetic features between reference and other selected organisms.

Here *Paenarthrobacter aurescens* Strain NBRC 12136 shares the close similarity with *Arthrobacter* sp. ATA002 but comparatively distantly related with *Arthrobacter* sp. GN70. *Arthrobacter sp. L77, Arthrobacter agilis* strain UMCV2 are distantly related to *Paenarthrobacter aurescens* Strain NBRC 12136, *Arthrobacter* sp. ATA002, *Arthrobacter* sp. GN70, *Arthrobacter sp. SPG23, Arthrobacter* sp. AFG7.2, *Arthrobacter ramosus* ZX_2022b, *Arthrobacter bambusae* DP-A9, *Arthrobacter sp.* UKPF54-2, *Pseudarthrobacter psychrotolerans* YJ56. Under the common clade *Arthrobacter ramosus* ZX_2022b, *Arthrobacter bambusae* DP-A9, *Arthrobacter sp.* UKPF54-2, *Pseudarthrobacter psychrotolerans* YJ56 are distantly related with *Arthrobacter* sp. ATA002, *Arthrobacter* sp. GN70. *Paenarthrobacter* AB444 was found to be most similar with *Paenarthrobacter histidinolovorans* JCM, *Paenarthrobacter ilicis* DSM 20138, and *Paenarthrobacter nitroguajacolicus* 3945 than *Paenarthrobacter aurescens* Strain NBRC 12136. The closest association of *Paenarthrobacter* AB444 was found to be with *Paenarthrobacter histidinolovorans* JCM with sharing same clade. *Paenarthrobacter nicotinovorans* ATCC 49919 found to possess moderate dissimilarity with *Paenarthrobacter* AB444 under a different clade. Among the *Paenarthrobacter group* the highest distant connection was of *Paenarthrobacter* AB444 was found to be with *Paenarthrobacter ureafaciens* DnL1-1. *Pseudarthrobacter* sp shows more distance *Paenarthrobacter* AB444 with different shared clade.

In VIP score, the gradient of shades directs the association of the individual gene cluster with specific microbial groups. The alteration in color with the gradient for individual gene clusters proposes an intense association among microbial groups. This approach enables to rapidly get the gene clusters that are significant concerning maximum VIP Score and the considerable groups that are mostly associated with. Here the VIP score plot highlights the importance of NRP, NRP-associated genes and terpene in discriminating among the *Paenarthrobacter* sp*, Pseudarthrobacter* sp and *Arthrobacter* sp (Figure 8b). The Venn diagram in (Figure 8c) also demonstrates the overlapping genetic features between the reference organism *Paenarthrobacter* AB444 (Query organism) and other organisms (*Arthrobacter agilis* strain UMCV2, *Arthrobacter bambusae* DP-A9, *Arthrobacter koreensis* 5J12A, *Arthrobacter ramosus* ZX_2022b, *Arthrobacter* sp. AFG20, *Arthrobacter* sp. AFG7.2, *Arthrobacter* sp. ATA002, *Arthrobacter* sp. *Arthrobacter* EpRS66, sp. EpRS71, *Arthrobacter* sp. GN70, *Arthrobacter sp. L77, Arthrobacter sp. PGP41, Arthrobacter sp. SPG23, Arthrobacter sp.* UKPF54-2, *Arthrobacter wenxiniae* strain AETb3-4, *Paenarthrobacter aurescens* Strain NBRC 12136, *Paenarthrobacter histidinolovorans* JCM 2520, *Paenarthrobacter ilicis* DSM 20138, *Paenarthrobacter nicotinovorans* ATCC 49919, *Paenarthrobacter nitroguajacolicus* 3945, *Paenarthrobacter ureafaciens* DnL1-1, *Pseudarthrobacter chlorophenolicus* A6, *Pseudarthrobacter chlorophenolicus* strain Mor30.16, *Pseudarthrobacter humi* RMG13, *Pseudarthrobacter psychrotolerans* YJ56, *Pseudarthrobacter siccitoleran*s Strain 4J27). The figure indicates that a total of 2, 1, 3, 2,3,3,3 genes were overlapping with the reference organism and selected organisms. The name of the selected organisms is provided in each of these Venn diagrams. In Figure 9a, the unsupervised PCA plot showcases the principal components (PC1 and PC2) responsible for separating the groups. The x-axis corresponds to PC1, which accounts for 26.6% of the variance, while the y-axis represents PC2, explaining 17.1% of the variance. Figure 9b and 10 depict the supervised PLS-DA plot, which further emphasizes the separation between the groups, with PC1 contributing 71% and PC2 contributing 6% to the variance.

**Figure 9.**
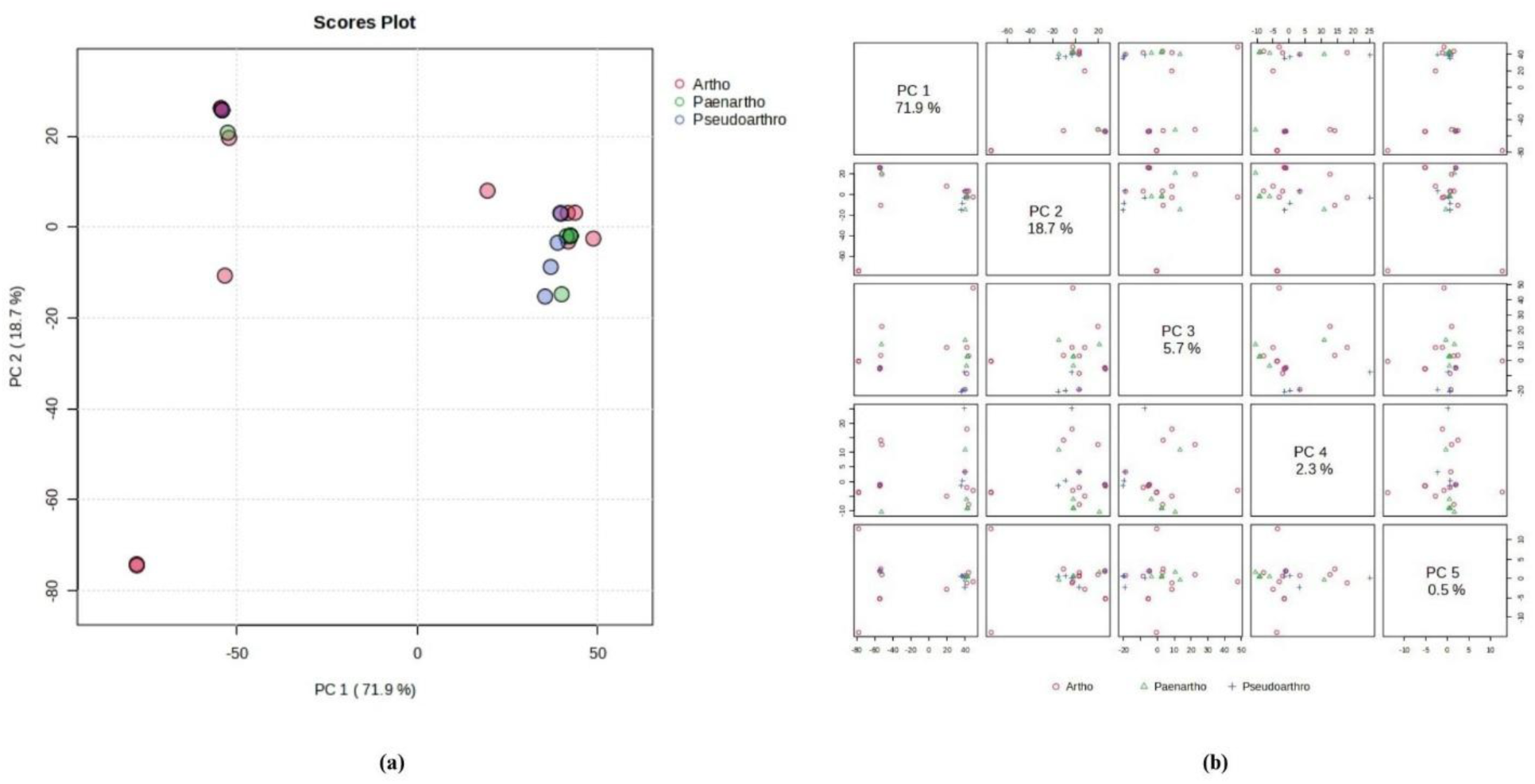
**(a).** The Unsupervised PCA plot showcasing the principal components (PC1 and PC2) responsible for separating the groups**, (b):** Different variances for each direction in PLS-DA plot

**Fig 10.**
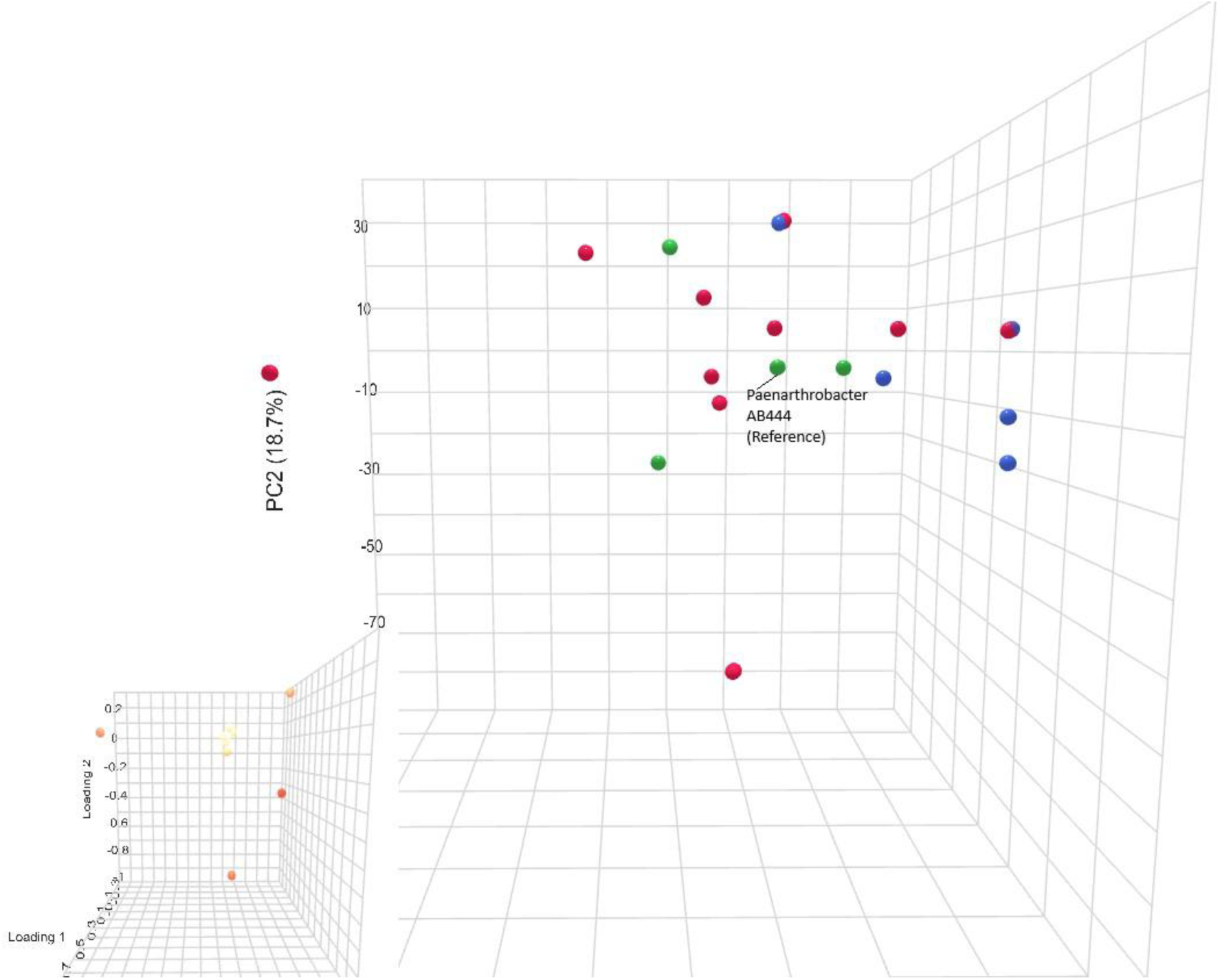
Supervised 3D PLS-DA emphasizing the separation between the groups, with PC1 and PC2

## 4. Discussion

Crop harvesting in the district of Darjeeling is hard to continue due to the acidic character of the soil. A number of fertilizers comprising nitrogen fertilizers also facilitate this acidic nature. This acidic nature of the soil ranges between (4.2 < pH < 7.0) that supports the replacement of a number of cations [sodium (Na), potassium (K), magnesium (Mg), calcium (Ca)] with protons along with a probability of leaching out of them from the rhizospheric zone [35]. Lower accessibility of these cations in combination with acidic nature of the soil hinders crop cultivation of this region significantly [36,37]. This agronomical perspective favours the practice of plant growth-promoting rhizobacteria by means of direct and indirect processes [6]. The rhizospheric zone of tea plant harbors assorted plant growth-promoting rhizobacteria that favor substantial use of them as sustainable biofertilizer within that agronomical system [38]. Hence, we go after the isolation, characterization, and study its potential along with its systemic study with closely related *Paenarthrobacter nicotinovorans*. The polyphasic detection of the selected rod-shaped microbe uncovered its biochemical pattern (S1 FILE), utilization profile of carbon source (S1 FILE), chemotaxonomic character. These patterns of a strain are likely to be operated by its exposure in the habitat. The metabolic edifice of the selected strain was established by partaking in the chemical and carbon source utilization assay. The Table S1 indicates that the strain prefers disaccharides more than other sugar derivatives such as N-Acetyl-β-D Mannosamine, β-Methyl-D Glucoside, 3-Methyl Glucose and most of sugar alcohols [39]. The growth of *Actinomyces viscosus* was inhibited in the presence of sugar alcohol and it could be a relatable one for the current strain [40]. [41] also suggests the involvement of repeating disaccharide units for building of microbial structural part. Mostly all amino acids were consumed effectively except D-Aspartic Acid. In *Arthrobacter protophormiae*. D-Aspartic Acid was also not taken as substrate [42]. It was found that naturally occurring myo-inositol can be used as carbon and energy source in *Arthrobacter globiformis* and it could be a supportive one for the characterization of the current strain [43]. Inosine was also consumed by the strain. Some of the antibiotics, such as Rifamycin SV, Nalidixic Acid and Troleandomycin were also used by the strain whereas Lincomycin and Minocycline showed borderline consumption [44]. Tetrazolium Blue and Tetrazolium Violet was consumed by the strain at a minimal level. Niaproof4 (surfactant) was not utilized by the strain and *Arthrobacter endophyticus* sp. nov., showed the same response [44]. Like other *Arthrobacter endophyticus,* sodium Butyrate the mammalian cell proliferation inhibitor was also used up by this strain [44].

FAME analysis process was found to be very helpful in recognizing and categorizing microbe on the basis of the unique profile of their fatty acid content. The spotted fatty acids and their comparative abundance are decisive in getting the cellular membrane constitution, environmental adaptability along with the establishment of phylogenetic relationship. 14:0 ISO, a branched fatty acid chain mostly located in actinomycetes, provides membrane stability to the strain. 15:0 ISO (5.33 %) is an iso-branched fatty acid chain identified to effect membrane fluidity of strain and is distinctive in actinomycetes. 15:0 ANTEISO A (49.07 %), defines the presence of the Anteiso branched fatty acid chain in the strain supporting the adaptation of the strain at a varied range of temperatures. 15:1 ANTEISO A (7.23 %) is a mono-unsaturated branched fatty acid chain having an anteiso pattern of branching. This double offers the membrane fluidity though the anteiso pattern maintains ideal membrane activity at low temperature. 16:0 ISO (19.69%) is a substantial fatty acid component of the strain influencing its membrane construction. 16:0 (2.43%) is a frequently found straight-chain fatty acid in bacterial membranes. The presence of 17:0 ANTEISO (10.87%) offers a thermal adaptation to the strain by associating with flexibility in low temperatures. Anteiso patterns of fatty acids are found to be accumulated in organisms subjected to lower temperatures. The abundance of iso, as well as anteiso pattern of branched fatty acids, is a typical character of the actinomycete group. Specific pathways that are followed within the microbial system for the synthesis of these fatty acids signify the metabolic and genetic capabilities of the microbes. Hence the profile of fatty acids aligns with the identified arrays in actinomycetes that characteristically show a high percentage of branched fatty acid chains. It proposes that the microbial strain is a part of this group. 15:1 ANTEISO A is a distinctive chemotaxonomic indicator, mostly related to particular actinomyces species. Chemotaxonomic profiling was consistent with the adherent of the genus *Paenarthrobacter* [45]. The observed phylogenetic clustering of *P. nicotinovorans* strains in the phylogenetic tree, combined with the pangenomic analysis, suggests a correlation between genetic relatedness and the presence of specific genomic traits. The conserved features within certain clades may represent core genes critical for the ecological niche or metabolic functions of these strains. In contrast, the sporadic distribution of other features may indicate adaptive responses to varying environmental conditions, such as different substrates or stressors.

The phylogenetic clustering of strains like AB444, *P. nicotinovorans* NPDC089708 and DS1097 suggests a shared evolutionary history, possibly shaped by similar selective pressures. These findings emphasise the role of genomic plasticity in the ecological success of *P. nicotinovorans*. Further studies integrating functional assays with genomic data could provide deeper insights into the specific roles of these genomic features in plant growth promotion, or other environmental adaptations. Draft genome sequence of *Peanarthrobacter* sp. AB444 revealed the presence of the essential genes involved in plant growth promotion activity. The function of the core genes related to phosphate solubilization, indole acetic acid (IAA) production, siderophore production, biofilm formation is reflected in its phenotypic characters as well. Genome analysis of AB444 revealed that like *Paenibacillus sonchi,* AB444 contains high affinity phosphate-specific transport (Pst) operon comprising of four genes, vis. *pstS*, *pstC*, *pstA* and *pstB* [46]. Beyond that presence of alkaline phosphatase, phosphodiesterase, phospholipase enable AB444 to solubilize complex inorganic and organic phosphates [47]. Its inorganic phosphate solubilization ability comes from the presence of inorganic pyrophosphatase (EC 3.6.1.1), exo-polyphosphatase (EC 3.6.1.11), phosphatase kinase (EC 2.7.4.1) and phosphate transporter genes [48,49]. AB444 contains a set of genes involved in the synthesis of bioactive enzymes like amylases, lipases, protease, chitinase which can be beneficial to inhibit the growth of bacterial and fungal phytopathogens. Riboflavin is an important component in plant defense. It functions as either elicitor or protectant molecule, so riboflavin producing PGPR might initiate the induced systemic defense responses in plants [50]. AB444 also contains a riboflavin synthase gene which might be beneficial in inducing systemic response in the host plant, keeping it aware of external stressed conditions [50]. Quorum sensing (QS) is an important phenomenon of PGPR. It helps them to survive in the rhizosphere by maintaining their population according to the environmental conditions [51]. LuxR is one of the important gene family of QS mechanism which functions as a transcriptional regulator. The draft genome indicates the presence of LuxR family genes in *Peanarthrobacter* sp. AB444 involved in two component signalling system. This finding aligns with the genome of another PGPR *Arthrobacter* sp. UMCV2 [51]. It has a wide range of genes involved in heavy metal degradation machinery, like copper uptake protein (YcnI), magnesium and cobalt efflux protein (CroC), Cobalt/zinc/cadmium resistance protein (CzcD), Arsenate reductase (EC 1.20.4.4) etc. Application of chemical fertilizers often lead to accumulation of heavy metals in soil, so rhizospheric microbes are generally well-adapted to the heavy metal degradation process. Hence, it is not surprising that AB444 also contains the heavy metal degradation/resistance cassette. Further, it will be advantageous to utilize the isolate in the heavy metal polluted cultivation areas [52]. This isolate has glycerol uptake and G3P dehydrogenase which indicates its ability to metabolize glycerol. It has proline/ glycine betaine transporter, along with acetoin, butanediol biosynthesis mechanism. Its biofilm formation ability is also reflected by the presence of biofilm forming genomic island. It also harbours EPS producing genes, along with several antibiotic biosynthesis machinery like phenazine synthesis (*PhnZ*), spermidine synthesis etc.

Accessory and unique genes, however, provide insights into the adaptive potential and ecological versatility of these strains. The enrichment of accessory genes in pathways like membrane transport and signal transduction suggests their role in enabling the bacteria to respond to environmental changes or stresses. Similarly, the presence of unique genes in pathways related to secondary metabolite biosynthesis and drug resistance underscores the strain-specific adaptations, which may be crucial for survival in unique environmental niches. The enrichment of pathways in the pangenome of AB444 and *P. nicotinovorans* strains related to xenobiotics biodegradation and environmental adaptation within the accessory and unique gene clusters of *P. nicotinovorans* underscores its potential as a plant growth-promoting rhizobacterium (PGPR). These pathways suggest that *P. nicotinovorans* is well-equipped to survive in diverse and challenging environments, including those contaminated with pollutants. Its ability to degrade xenobiotic compounds not only aids in soil detoxification but also creates a more favorable environment for plant growth. Furthermore, the presence of genes involved in nutrient cycling, such as those related to nitrogen and phosphate metabolism, positions AB444 as a candidate for promoting sustainable agriculture. These traits can enhance nutrient availability and uptake by plants, fostering improved growth and productivity. The pathways associated with environmental adaptation further indicate that AB444 can thrive under abiotic stress conditions, such as salinity or drought, which are critical for supporting plant resilience in adverse environments. Overall, the functional diversity of the accessory and unique gene clusters highlights AB444 as a promising PGPR with applications in bioremediation and sustainable agriculture, enabling it to play a dual role in improving soil health and enhancing plant growth.

Among the regions of the biosynthetic gene cluster, one region with highest similarity (100%) to known cluster, that is related to the desferrioxamine E biosynthesis; another one with less similarity (20%-40%), those related to the carotenoid and stenothricin biosynthesis. Three regions such as Type III Polyketide synthase, betalactone and amglyccycl with very low similarity (6%-7%) with the known clusters, those related to the SW-163C, UK-63598, SW-163E, SW-163F, SW-163G, microansamycin, cetoniacytone biosynthesis. Comprehensive analysis of multiple biosynthetic genes of functionally related microbial counts in our study revealed that *Paenarthrobacter* sp. AB444 has vibrant evolutionary pattern like horizontal gene transfer (HGT), gene duplication and molecular reorganizations alike other strains of Paenarthrobacter and Arthrobacter genera. The *Paenarthrobacter* sp. AB444 shows a high rate of horizontal gene transfer and duplication like *Arthrobacter* sp. ATA002. It connects to the dynamicity of the production of metabolites [53]. Strains such as *Arthrobacter agilis* UMCV2 and *Pseudarthrobacter siccitolerans* both of them are advanced by means of reorganization and additions of genes. Still, it covers a very narrower diversity of BSG that counts for their novel product formation limitation [14,54]. Strains such as *Arthrobacter koreensis* 5J12A and *Arthrobacter* sp. ATA002 are capable in production of enhanced polyketides and nonribosomal peptides. Whereas, *Paenarthrobacter* sp. AB444 was found to be more focused on the production of metabolites responsible for the antimicrobial activities and stress response, as implied from the genomic closeness to the strains that are capable of bioremediation like *Paenarthrobacter nitroguajacolicus*[14]. *Paenarthrobacter ilicis* DSM 20138 and *Peanarthrobacter* sp. AB444 carries BSGs that are capable of producing antifungal products. The variety of molecular PKS genes of *Peanarthrobacter* sp. AB444 leave behind the related strain revealing this strains as potential source for the new antibiotic synthesis. The coherent difference of the *Arthrobacter/Pseudoarthrobacter/Paenarthrobacter* groups in three individual clades is much favoured by the gene support indices (GSIs)[14]. The relative study of *Paenarthrobacter* sp. AB444 with closely related stains underlines the implication of the variety of BGCs in microbial flexibility and metabolite biosynthesis [55]. Overall, these visualizations provide insights into the relationships between species and genes, as well as the discriminatory power of different variables in distinguishing between groups. This establishes the significant difference between *Paenarthrobacter* sp. AB444 and other *Arthrobacter* sp, *Pseudoarthrobacter* sp.

## 5. Conclusion

These analyses offer a detailed understanding of the metabolic potential and ecological strategies utilized by the AB444 strain, highlighting its capabilities as a plant growth-promoting rhizobacterium (PGPR). The identified pathways feature its adaptability to diverse environments and its potential role in enhancing plant growth and resilience. Future research combining functional assays with genomic insights could provide further validation of these findings, paving the way for novel applications of the AB444 strain in sustainable agriculture and environmental management. Such studies could also uncover additional mechanisms by which AB444 contributes to plant health and soil fertility, reinforcing its value as a multifunctional PGPR.

## Author contributions

Triparna Mukherjee, Conceptualization, Data curation, Formal analysis, Investigation, Methodology, Validation, Visualization, Writing – original draft, Writing – review and editing, Funding acquisition | Sangita Mondal, Formal analysis, Investigation, Methodology| Dhruba Bhattacharya, Investigation, Methodology | Sanjukta Dasgupta, Formal analysis, Methodology| Abhrajyoti Ghosh, Conceptualization, Formal analysis, Funding acquisition, Investigation, Methodology, Project administration, Resources, Supervision, Writing – original draft, Writing – review and editing

## Acknowledgements

This work was supported by the ICMR Research Associate scheme (File no.45/18/2019-PHA/BMS ) by the Indian Council of Medical Research (ICMR), Government of India and TM was supported by this ICMR Research Associateship. SM was supported by a Senior Research Scholarship (Ref. 191620161234) from the University Grant Commission (UGC), India. DB was supported by SERB-National Postdoctoral Fellowship (SERB-NPDF) [Ref. PDF/2021/003080] from SERB, India. The authors would like to acknowledge the Central Instrument Facility (CIF) of Bose Institute, India.

## Data availability

The raw draft genome sequence and assembled genome of AB444 obtained from this study have been deposited in the NCBI under a single bioproject with the accession number PRJNA931222.

---

**Figure S1.**
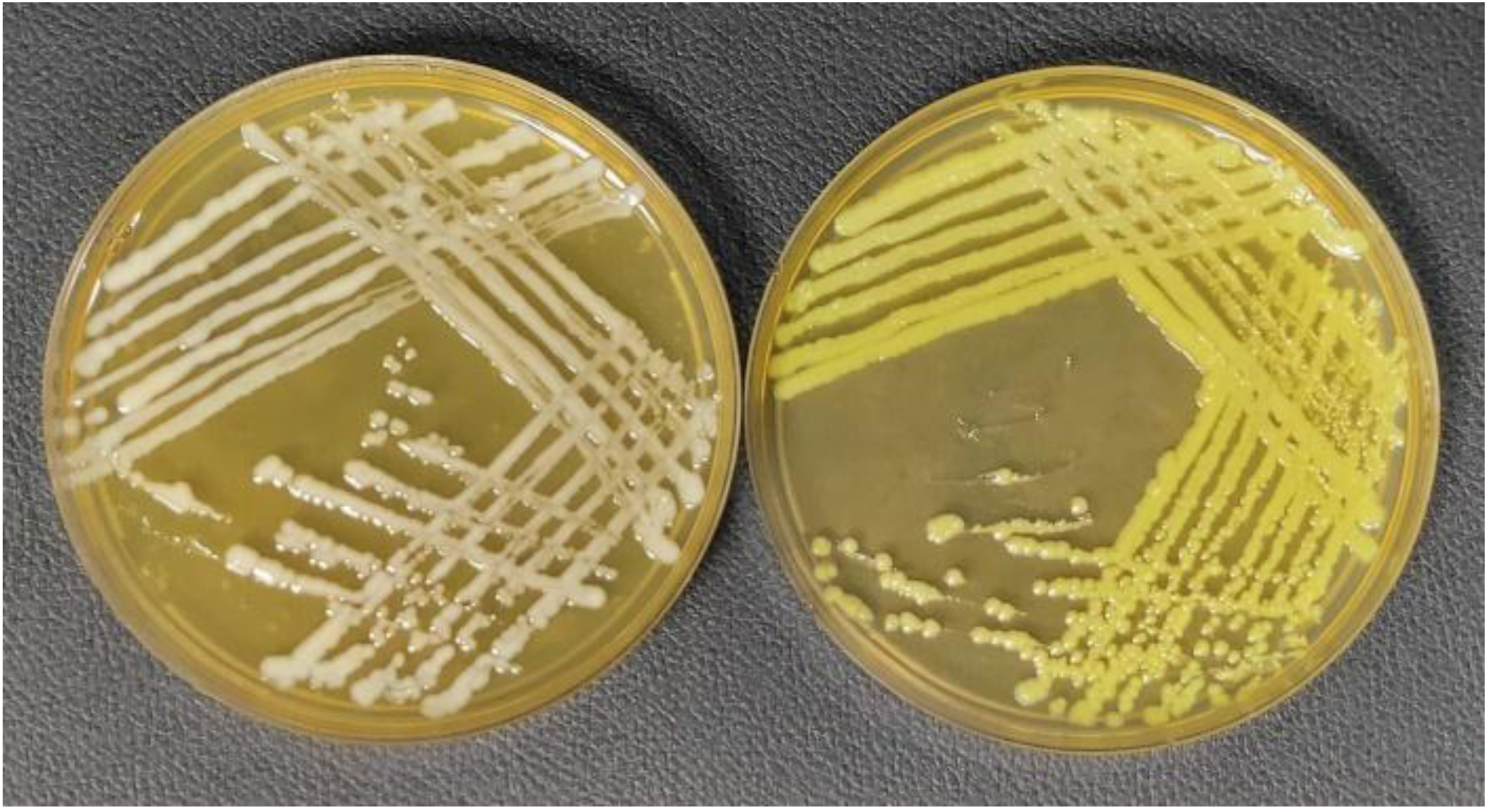
Colonies grown under light conditions appear vivid or pale yellow, whereas those grown under dark conditions appear off-white or cream.

**Figure S2.**
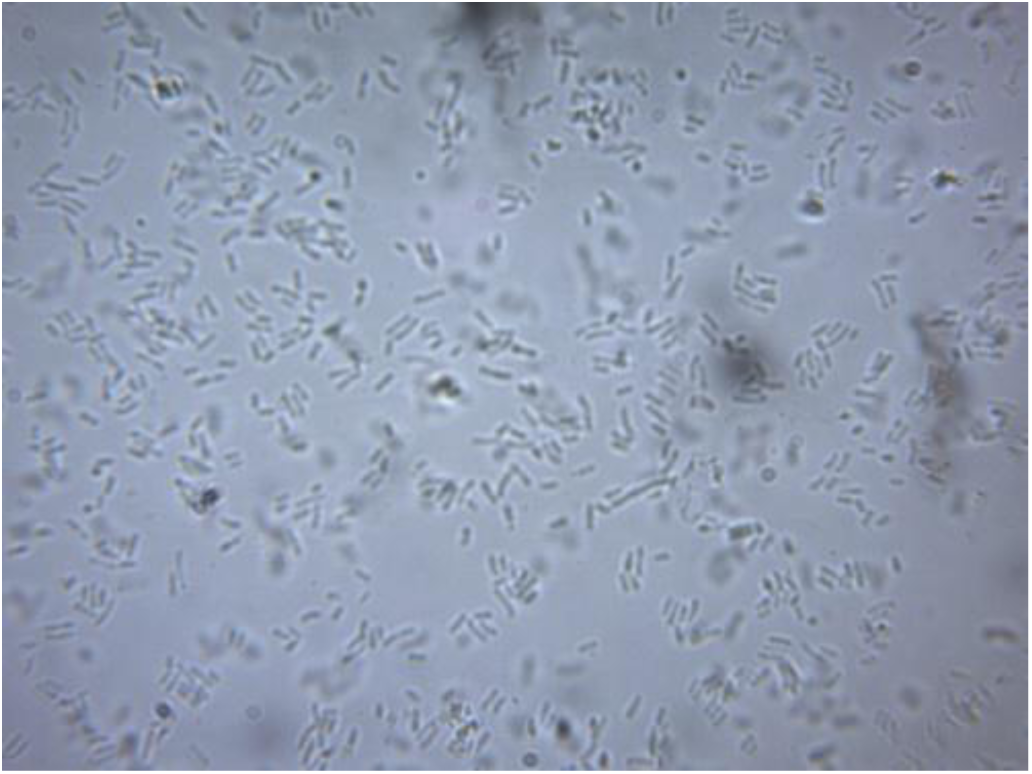
Arrangement of mycelium studied by phase contrast microscopy. Phase contrast micrographs of actinomycets *Paenarthrobacter* sp. Cells were found as rod-shaped with some cell forming v-shaped arrangement. They were found to be motile in nature.

**Figure S3:**
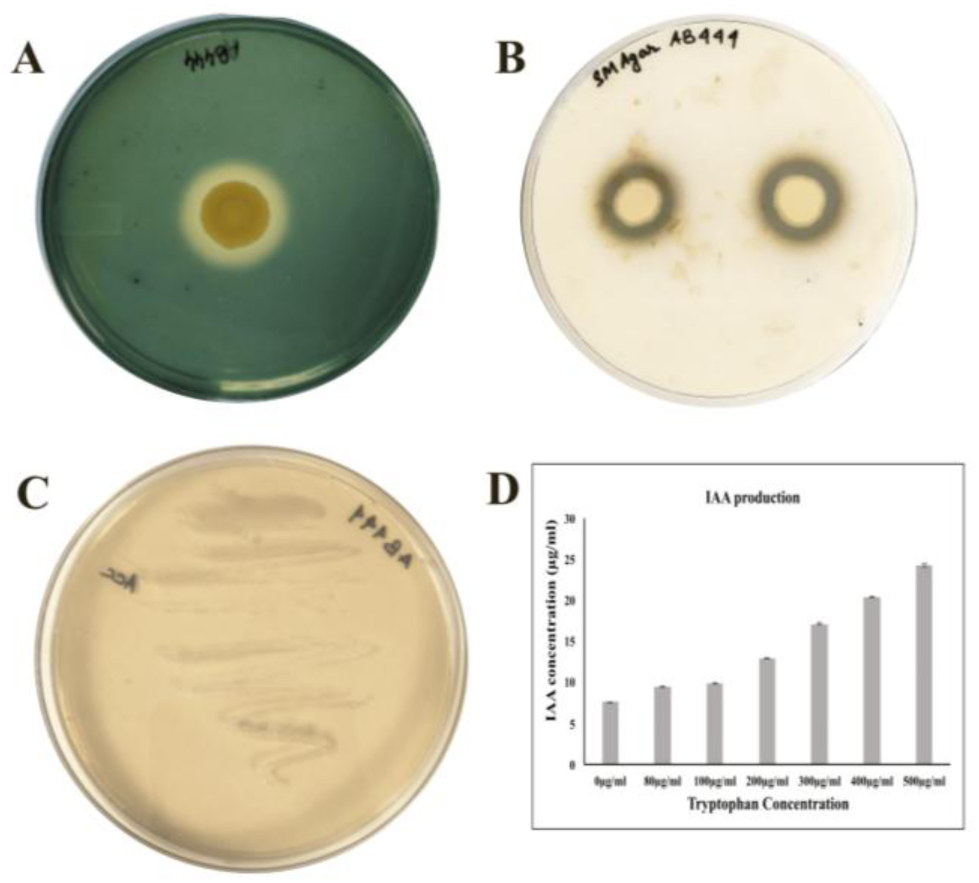
3A. Siderophore, 3B. protease, 3C. IAA and 3D. ACC deaminase

**Figure S4:**
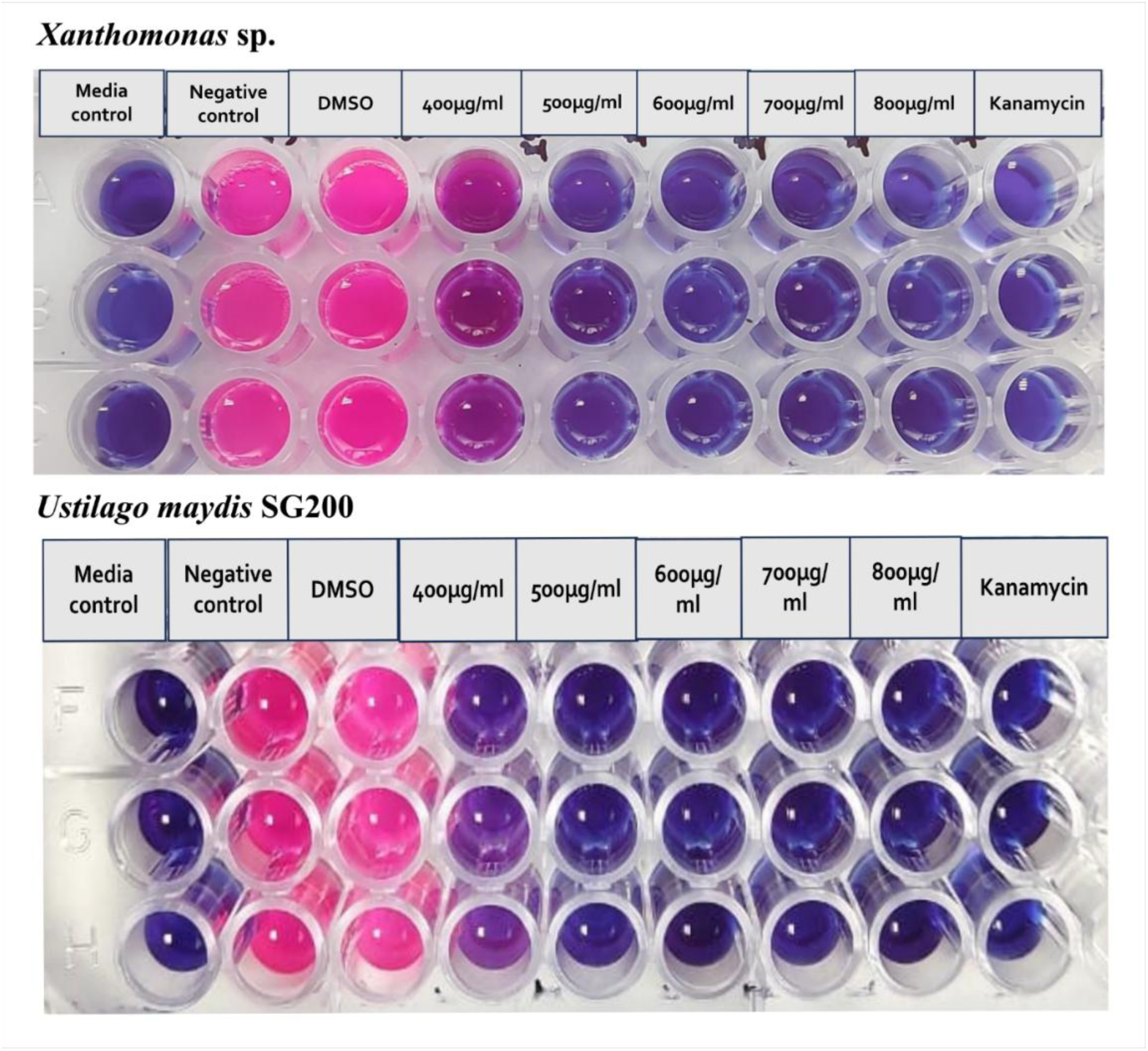
Phytopathogenic activity of AB444

**Figure S5:**
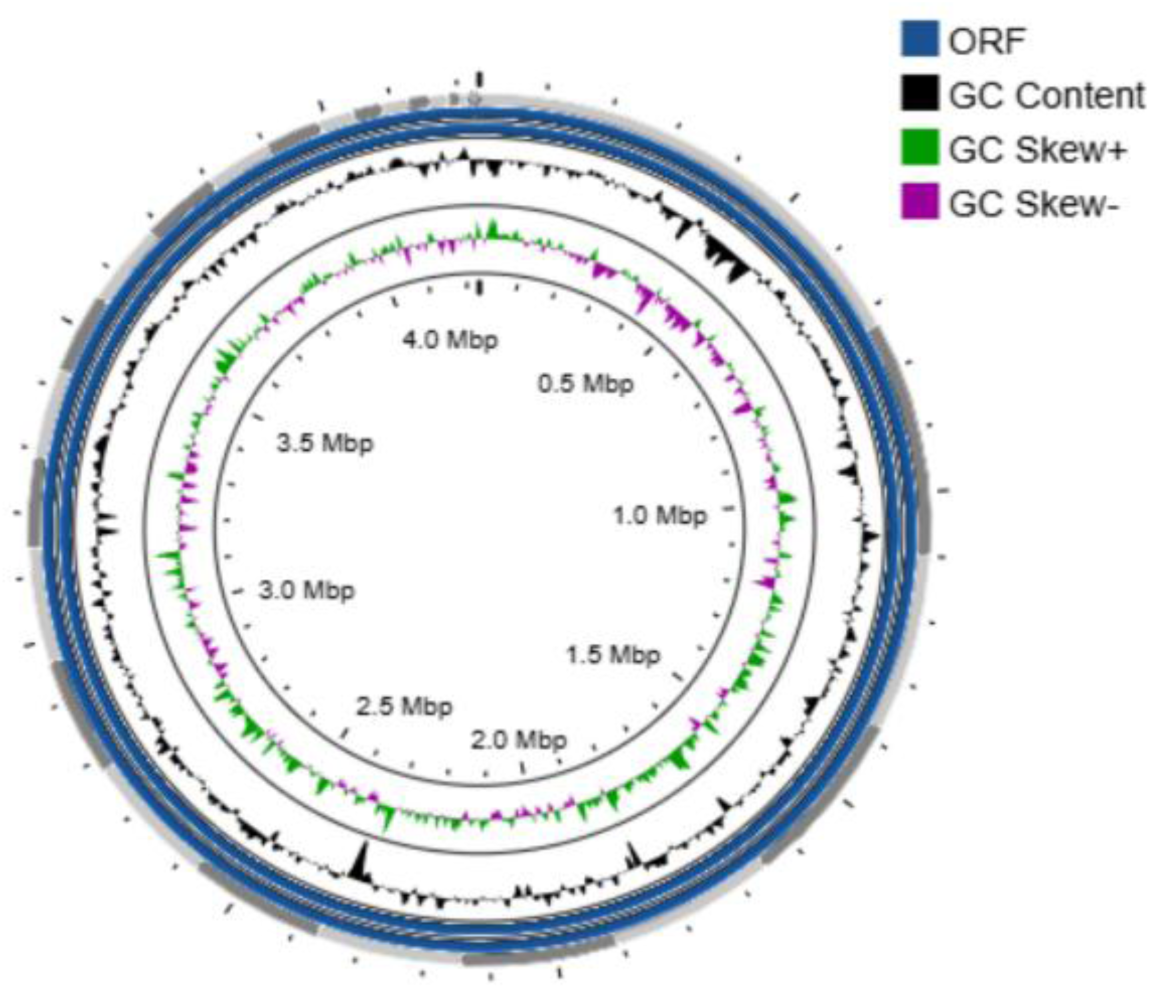
Circular genome map of AB444 showing GC content, GC skew

**Figure S6:**
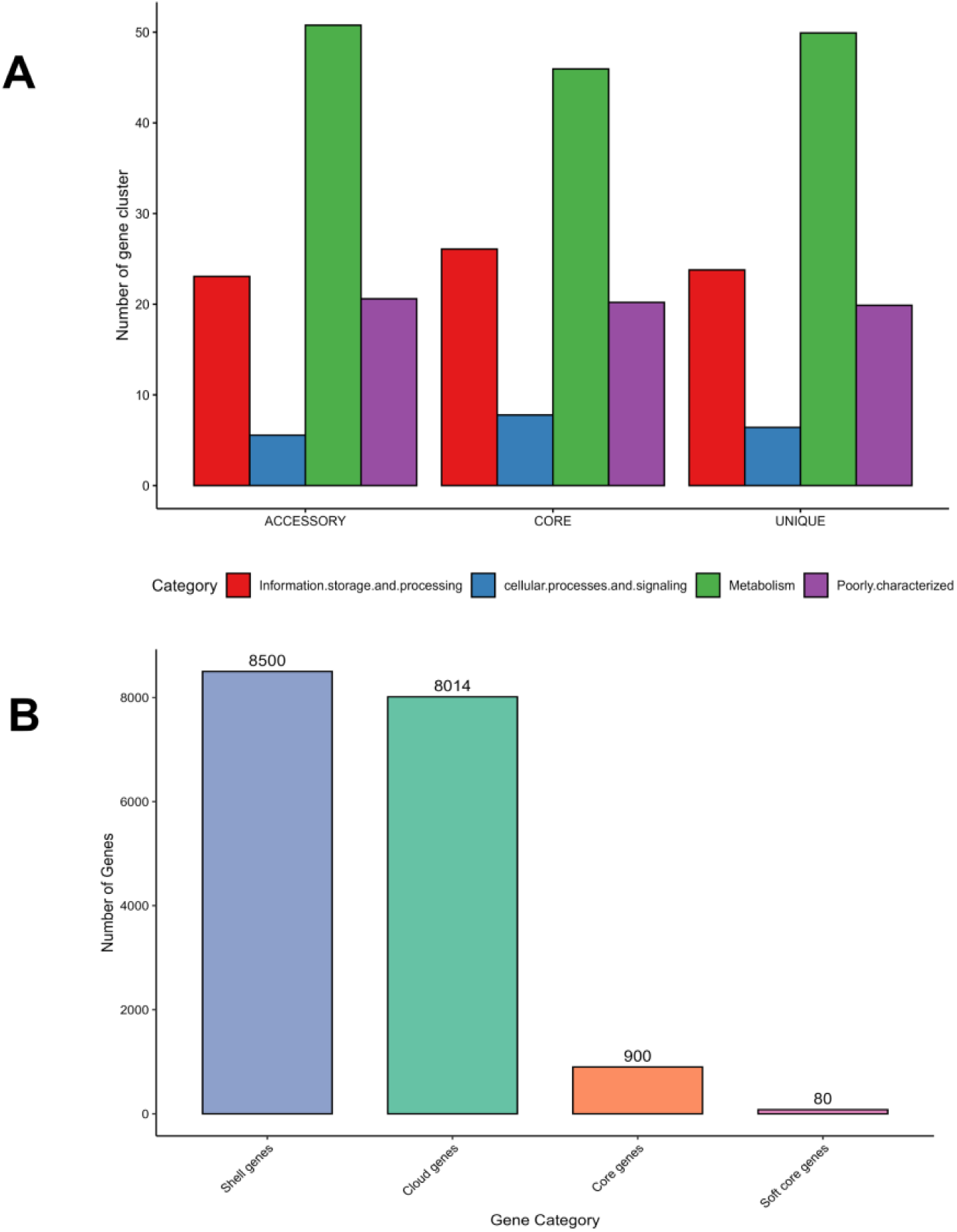
Number of gene clusters and categories obtained from pangenomic analysis of AB444 and *P. nicotinovorans* genomes

**Table S1:** Resource utilization pattern using Biolog

| Name of TEST | Result | Name of TEST | Result |
| --- | --- | --- | --- |
| Negative Control | Negative | L-Fucose | Borderline |
| Dextrin | Borderline | L-Rhamnose | Borderline |
| D-Maltose | Positive | Inosine | Positive |
| D-Trehalose | Positive | 1% Sodium | Borderline |
| D-Cellobiose | Positive | Fusidic Acid | Negative |
| Gentiobiose | Positive | D-Serine | Positive |
| Sucrose | Positive | D-Sorbitol | Negative |
| D-Turanose | Positive | D-Mannitol | Positive |
| Stachyose | Positive | Lactate | Nil |
| Positive Control | Positive | D-Arabitol | Negative |
| pH 6 | Borderline | myo-Inositol | Positive |
| pH 5 | Positive | Glycerol | Positive |
| D-Raffinose | Positive | D-Glucose 6-PO4 | Negative |
| α-D-Lactose | Mismatch<br>Positive | D-Fructose 6-PO4 | Borderline |
| D-Melibiose | Positive | D-Aspartic Acid | Negative |
| **β-Methyl-D Glucoside** | **Borderline** | **Troleandomycin** | **Borderline** |
| **D-Salicin** | **Positive** |  |  |
| **N-Acetyl-D Glucosamine** | **Positive** | **Rifamycin SV** | **Positive** |
| **N-Acetyl-β-D Mannosamine** | **Negative** | **Minocycline** | **Borderline** |
| **N-Acetyl-D Galactosamine** | **Mismatch Positive** | **Gelatin** | **Positive** |
| **N-Acetyl Neuraminic Acid** | **Positive** | **Glycyl-L-Proline** | **Positive** |
| **1% NaCl** | **Positive** | **L-Alanine** | **Positive** |
| **4% NaCl** | **Positive** | **L-Arginine** | **Borderline** |
| **8% NaCl** | **Borderline** | **L-Aspartic Acid** | **Positive** |
| **α-D-Glucose** | **Positive** | **L-Glutamic Acid** | **Positive** |
| **D-Mannose** | **Positive** | **L-Histidine** | **Positive** |
| **D-Fructose** | **Positive** | **L-Pyroglutamic Acid** | **Positive** |
| **D-Galactose** | **Positive** | **L-Serine** | **Positive** |
| **3-Methyl Glucose** | **Negative** | **Lincomycin** | **Borderline** |
| **D-Fucose** | **Negative** | **Guanidine HCl** | **Positive** |
| **Pectin** | **Borderline** | **Niaproof 4** | **Negative** |
| **D-Galacturonic Acid** | **Mismatch positive** | **Mucic Acid** | **Negative** |
| **L-Galactonic Acid Lactone** | **Negative** | **D-Saccharic Acid** | **Negative** |
| **D-Gluconic Acid** | **Positive** | **Vancomycin** | **Mismatched positive** |
| **D-Glucuronic Acid** | **Positive** | **Tetrazolium Violet** | **Borderline** |
| **Glucuronamide** | **Baseline** | **Tetrazolium Blue** | **Borderline** |
| **p-Hydroxy Phenylacetic Acid** | **Borderline** | **Potassium Tellurite** | **Positive** |
| **Methyl Pyruvate** | **Borderline** | **Tween 40** | **Borderline** |
| **D-Lactic Acid Methyl Ester** | **Negative** | **γ-Amino-Butyric Acid** | **Borderline** |
| **Sodium Bromate** | **Positive** | **α-Hydroxy Butyric Acid** | **Borderline** |
| **L-Lactic Acid** | **Positive** | **β-Hydroxy-D,L Butyric Acid** | **Positive** |
| **Citric Acid** | **Positive** | **α-Keto-Butyric Acid** | **Borderline** |
| **α-Keto-Glutaric Acid** | **Mismatch positive** | **Acetoacetic Acid** | **Borderline** |
| **D-Malic Acid** | **Borderline** | **Propionic Acid** | **Positive** |
| **L-Malic Acid** | **Positive** | **Acetic Acid** | **Positive** |
| **Bromo-Succinic Acid** | **Positive** | **Formic Acid** | **Borderline** |
| **Nalidixic Acid** | **Positive** | **Aztreonam** | **Positive** |

**Table S2:**
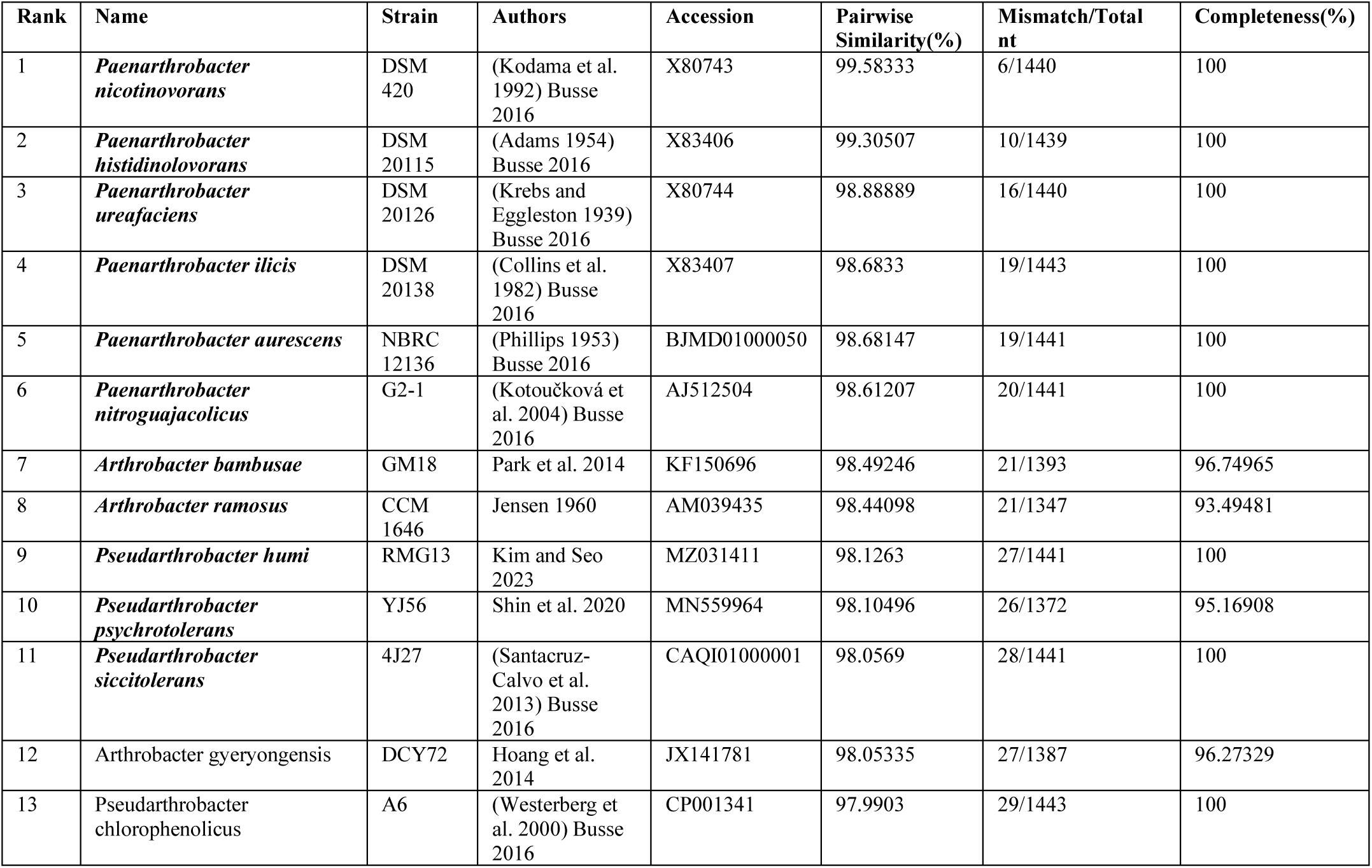
List of bacterial strains showing 16S rRNA homology with AB444

**Table S3:** *Paenarthrobacter nicotinovorans* genomes and their ANIb and DDH values calculated against AB444

| Assembly | GenBank | RefSeq | Genome | ANI% with AB444 | DDH % with AB444 |
| --- | --- | --- | --- | --- | --- |
| ASM2191934v1 | GCA_021919345.1 | GCF_021919345.1 | *Paenarthrobacter nicotinovorans* ATCC 49919 (strain) | 84.50 | 68.1 |
| ASM4352004v1 | GCA_043520045.1 | GCF_043520045.1 | *Paenarthrobacter nicotinovorans* WA1 (strain) | 84.48 | 69.3 |
| ASM2191800v1 | GCA_021918005.1 | GCF_021918005.1 | *Paenarthrobacter nicotinovorans* nic- (strain) | 84.51 | 71.2 |
| ASM1787644v1 | GCA_017876445.1 | GCF_017876445.1 | *Paenarthrobacter nicotinovorans* DSM 420 (strain) | 84.51 | 68.4 |
| ASM1464873v1 | GCA_014648735.1 | GCF_014648735.1 | *Paenarthrobacter nicotinovorans* JCM 3874 (strain) | 84.50 | 68.4 |
| ASM3988921v1 | GCA_039889215.1 | GCF_039889215.1 | *Paenarthrobacter nicotinovorans* Se16.17 (strain) | 81.14 | 45.5 |
| ASM3145457v1 | GCA_031454575.1 | GCF_031454575.1 | *Paenarthrobacter nicotinovorans* DS2026 (strain) | 81.05 | 45.8 |
| ASM767025v1 | GCA_007670255.1 | GCF_007670255.1 | *Paenarthrobacter nicotinovorans* SSBW5 (strain) | 97.49 | 83.9 |
| ASM4474906v1 | GCA_044749065.1 | GCF_044749065.1 | *Paenarthrobacter nicotinovorans* NPDC089694 (strain) | 98.35 | 90.8 |
| ASM3081255v1 | GCA_030812555.1 | GCF_030812555.1 | *Paenarthrobacter nicotinovorans* CC523 (strain) | 84.41 | 63.3 |
| ASM4474910v1 | GCA_044749105.1 | GCF_044749105.1 | *Paenarthrobacter nicotinovorans* NPDC089692 (strain) | 98.25 | 90.8 |
| ASM3081174v1 | GCA_030811745.1 | GCF_030811745.1 | *Paenarthrobacter nicotinovorans* DS1097 (strain) | 98.25 | 88.7 |
| ASM166244v1 | GCA_001662445.1 | GCF_001662445.1 | *Paenarthrobacter nicotinovorans* Hce-1 (strain) | 84.45 | 64 |
| ASM4474892v1 | GCA_044748925.1 | GCF_044748925.1 | *Paenarthrobacter nicotinovorans* NPDC089711 (strain) | 98.29 | 89 |
| ASM4474898v1 | GCA_044748985.1 | GCF_044748985.1 | *Paenarthrobacter nicotinovorans* NPDC089708 (strain) | 98.04 | 86.9 |
| ASM2995246v1 | GCA_029952465.1 | GCF_029952465.1 | *Paenarthrobacter nicotinovorans* LG02 (strain) | 84.71 | 56.3 |
| ASM51401v1 | GCA_000514015.1 | GCF_000514015.1 | *Paenarthrobacter nicotinovorans* 231Sha2.1M6 (strain) | 81.13 | 45.3 |
| ASM51415v1 | GCA_000514155.1 | GCF_000514155.1 | *Paenarthrobacter nicotinovorans* 26Cvi1.1E (strain) | 81.15 | 46.9 |
| ASM2416867v1 | GCA_024168675.1 | _____ | *Paenarthrobacter nicotinovorans* GSJI39 (strain) | 84.31 | 67.1 |

